# Spontaneous whole-genome duplication restores fertility in interspecific hybrids

**DOI:** 10.1101/538298

**Authors:** Guillaume Charron, Souhir Marsit, Mathieu Hénault, Hélène Martin, Christian R. Landry

**Author notes:** equal contributions from these authors (Guillaume Charron, Souhir Marsit).

## Abstract

Interspecies hybrids often show advantages over parents but suffer from reduced fertility, which can sometimes be overcome through sexual reproduction that sorts out incompatibilities. Sex is however inefficient due to the low viability or fertility of hybrid offspring and thus limits their evolutionary potential. Mitotic cell division could be an alternative to fertility recovery in facultative sexual species. To test this, we evolved under relaxed selection more than 600 diploid yeast hybrids between species that span 100,000 to 15 M years of divergence. We find that hybrids can recover fertility spontaneously and rapidly through whole-genome duplication. These events occurred in both hybrids between young and well-established species. Our results show that the instability of hybrid ploidy is a spontaneous path to fertility recovery.

**One Sentence Summary:** Ploidy changes potentiate hybrid speciation by leading to fertility recovery.

## Main Text

Inter-specific hybridization is common in animals, plants and microorganisms (*1, 2*) and is a potentially frequent source of genetic diversity over short time scales (*3, 4*). However, hybrid lineages often suffer from poor fertility that reflect reproductive isolation between parental lineages, which can hinder their potential as new species or populations. Different molecular mechanisms underly hybrid infertility, including genetic incompatibilities (nuclear and cytonuclear) and changes in genome architecture (ploidy number or chromosome rearrangements) (*5, 6*). If the hybrids are to establish as species, they need to recover from this low initial fitness by restoration of their fertility. In obligatory sexual species, fertility restoration can be achieved by crosses among hybrids or backcrosses with either parental species, allowing the purge of incompatibility through recombination, leading to the formation of introgressed species (*7*). Some organisms, however, have access to both sexual and asexual reproduction. In these species, if sexual encounters are rare, hybrids might be able to recover fertility by other means than recombination. As an example, somatic chromosome doubling in diploid tissues or zygotes can lead to the emergence of polyploids, which may display both restored fertility and reproductive isolation with parental species (*8*). Polyploidy is most common in plants (*9*) but has also been observed in animals and fungi (*10, 11*). Among the main questions that remain to be answered is how frequent fertility restoration without sexual reproduction is, what are the mechanisms by which fertility restoration occurs and whether it occurs without the intervention of natural selection.

We investigated the evolution of fertility in experimental yeast hybrids during mitotic evolution under strong population bottlenecking that minimizes the efficiency of selection. We examined whether or not fertility would increase or decrease with time and, if so, whether it would occur through gradual or punctuated changes (Figure 1A). We considered hybridization over a gradient of parental divergence from intra-population to inter-specific crosses, spanning up to 15 M years of divergence, which is sufficient to cause 99% reproductive isolation in budding yeast. To do so, we used a collection of North American natural yeast isolates representing three lineages of the wild species *Saccharomyces paradoxus* and a bona fide sister species, *S. cerevisiae* (Table S1). The *S. paradoxus* lineages (*SpA*, *SpB* and *SpC*)(*12*) are incipient species that exhibit up to 4% genetic divergence and up to 60% reduction of fertility in hybrids. They occur in partially overlapping geographical ranges, even for the most distant pair, *S. cerevisiae* and *S. paradoxus*, making hybridization possible. Including this sister species extends the genetic divergence of the crosses to 15%. For two independent strains per lineage, we mated *SpB* strains to other strains over a gradient of divergence (Very Low (VL_div_) = *SpB*×*SpB*, Low (L_div_) = *SpB*×*SpC*, Moderate (M_div_) = *SpB*×*SpA* and High (H_div_) = *SpB*×*S. cerevisiae*, Table S2, S3). Ninety-six independent diploid hybrid lines were generated for all but the VL_div_ crosses, for which 48 hybrids were generated, for a total of 672 lines (96 lines × 3 types of crosses × 2 pairs of strains, + 2×48). We randomly selected and streaked colonies on plates every 3 days for 35 passages (~22 mitotic divisions/passage) (Fig 1B, Fig. S1), which allowed to relax selection for growth and thus allowed genomic changes to fix randomly (*13*).

**Fig. 1.**
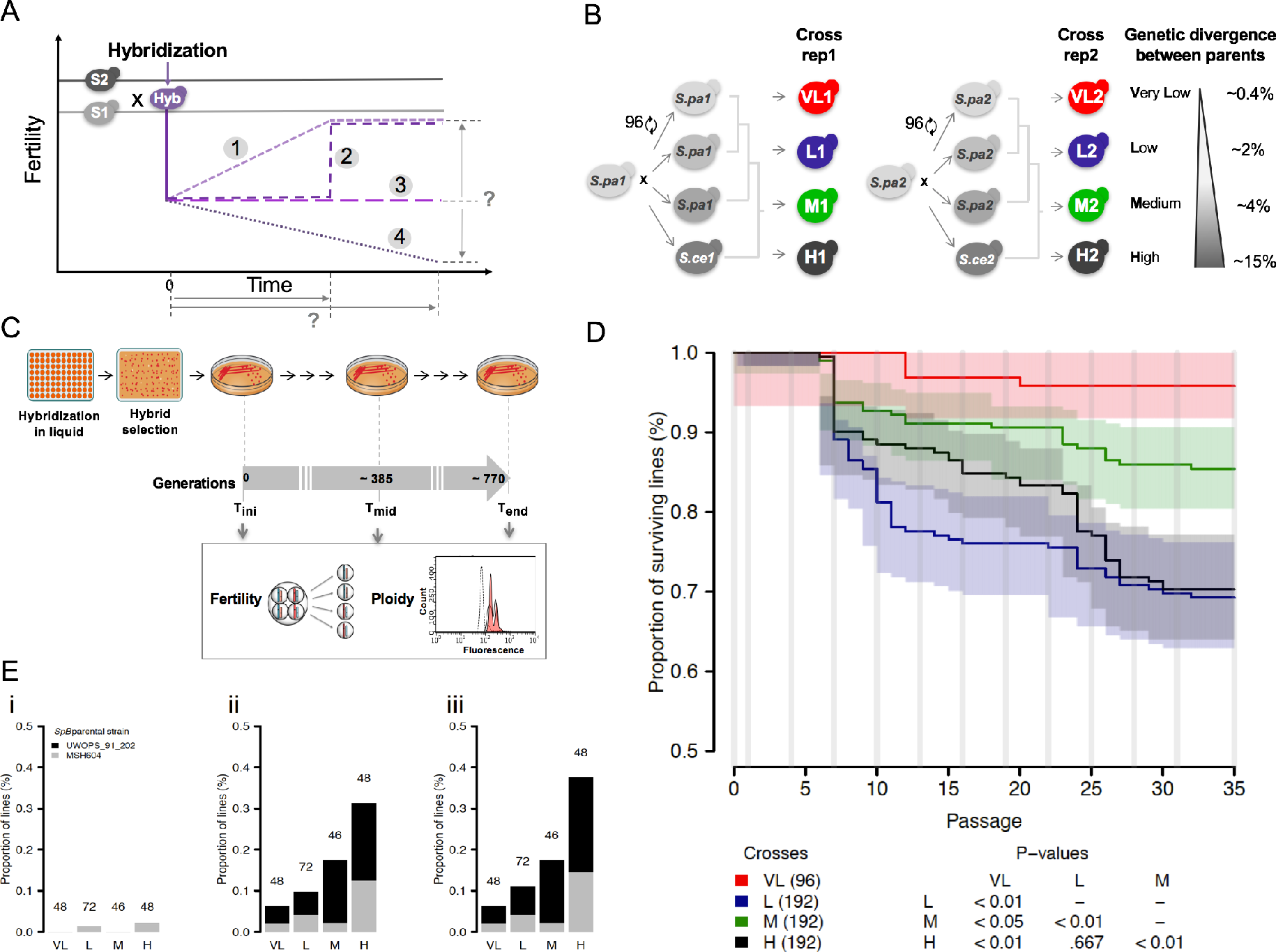
Neutral evolution of yeast hybrids shows the effect of genetic divergence on the evolution of viability and sporulation ability. (**A**) Evolution of fertility during mitotic proliferation of hybrids between species (S1 and S2): (1) Gradual recovery over time, (2) rapid recovery, (3) no significant recovery, (4) decline over time (**B**) Crosses were performed among *S. paradoxus* lineages (VL_div_, L_div_ and M_div_) and with *S. cerevisiae* (H_div_). (**C**) The 672 hybrids were evolved in conditions of weak selection to examine the neutral spontaneous evolution of fertility. Mitotic propagation was performed through repeated bottlenecks of single cells. Fertility was measured by estimating spore viability after meiosis and DNA content was measured by flow cytometry to estimate ploidy at T_ini_, T_mid_ and T_end_. (**D)**Survival rates vary among lines. Timing of each glycerol stock indicated by vertical grey lines. Logrank test pairwise comparisons FDR corrected P-values are shown. (E) Fraction of lines that lost their sporulation capacity at T_ini_ (i), T_mid_ (ii) and T_end_(iii). The number of strains tested per cross type is indicated over the corresponding bars.

After 770 mitotic generations, 77.9% of the lines (524 out of the 672 initial lines) were still able to grow. From the extinct lines, 48.6% (n= 72) were lost before 250 mitotic generations. This suggests that the loss of these lines is mostly due to genomic instability that arises rapidly after hybridization rather than spontaneous mutations, which would happen at a much slower pace (*14*). The L_div_ and H_div_ crosses had a significantly lower proportion of surviving lines (average of 69.3% and 70.3%, Fig. 1C, Fig. S2) compared to VL_div_ and M_div_ (averages of 95.8% and 85.4%, Fig. 1C, P < 0.01, Log-rank test, Table S4). This indicates that L_div_ and H_div_ hybrids may suffer from exacerbated genomic instability that lead to the rapid collapse of populations when faced with serial bottlenecks. This is in strong contrast with other observations reporting that yeast hybrids often show heterosis (*15, 16*) for growth performance and have access to more genetic variation to adapt to stressful conditions (*17*). Our results show that hybrids could also have access to more deleterious genomic changes whose effects are visible when allowed to randomly fix.

We measured fertility by inducing sporulation and meiosis and counting progeny survival in 214 randomly selected lines at three time points roughly corresponding to the initial (T_ini_, right after hybridization), middle (T_mid_, 385 generations) and terminal (T_end_, 770 generations) time points. Initial fertilities were consistent with previous estimates (< 1% for H_div_, 27.7% for M_div_, 34.2% for L_div_ and 60.3% for VL_div_, Fig. S3) (*18*). Unexpectedly, fertility could not be assessed for all the lines at T_end_ because 17% (n=37) of the tested lines lost their ability to enter meiosis (sporulation). Interestingly, the probability of sporulation is negatively correlated with parental divergence (Fig. 1D, r=−.76, P < 0.01, logistic regression). Sporulation requires functional aerobic respiration that requires the maintenance of functional mitochondrial DNA (mtDNA) (*19*), suggesting that loss of sporulation could be caused by the loss of mitochondrial functions. The genotyping of two mitochondrial loci revealed a strong association between loss of sporulation capacity and the absence of at least one mitochondrial marker and respiration (Supplementary text, Fig. S4A and S5, Fisher’s exact test, odds ratio > 77, P = 2.15×10^−6^). This suggests that mtDNA instability can contribute to reproductive isolation among closely related yeast populations and that this effect increases with genetic distance, as shown for more distant species (*20*).

To investigate whether fertility improved over the experiment, we calculated a fertility recovery score (FRS) as the difference in fertility between T_end_ and T_ini_ (Fig. 2A). As a point of comparison for fertility restoration with sexual reproduction, we performed 12 meiotic generations of intra-tetrad crosses (ITC) and calculated FRS, all of which were positive (Fig. 2B, Fig. S6). This is in stark contrast with mitotic lines, in which we found no bias in the change of fertility as the distribution of FRS was unimodal and centered on 0 (Fig. 2B, Fig. S7), showing that fertility is as likely to increase as it is to decrease. To make sure that low fertility was not due to the intrinsic inability of strains to produce viable spores, we performed autodiploidization on a random set of 16 haploid spores from the L_div_ and M_div_ crosses and this, at the three timepoints. In most cases, fertility was restored to more than 85% upon selfing (Fig. 2C), showing that infertility mostly derives from the presence of two divergent genomes in the same cell.

**Fig. 2.**
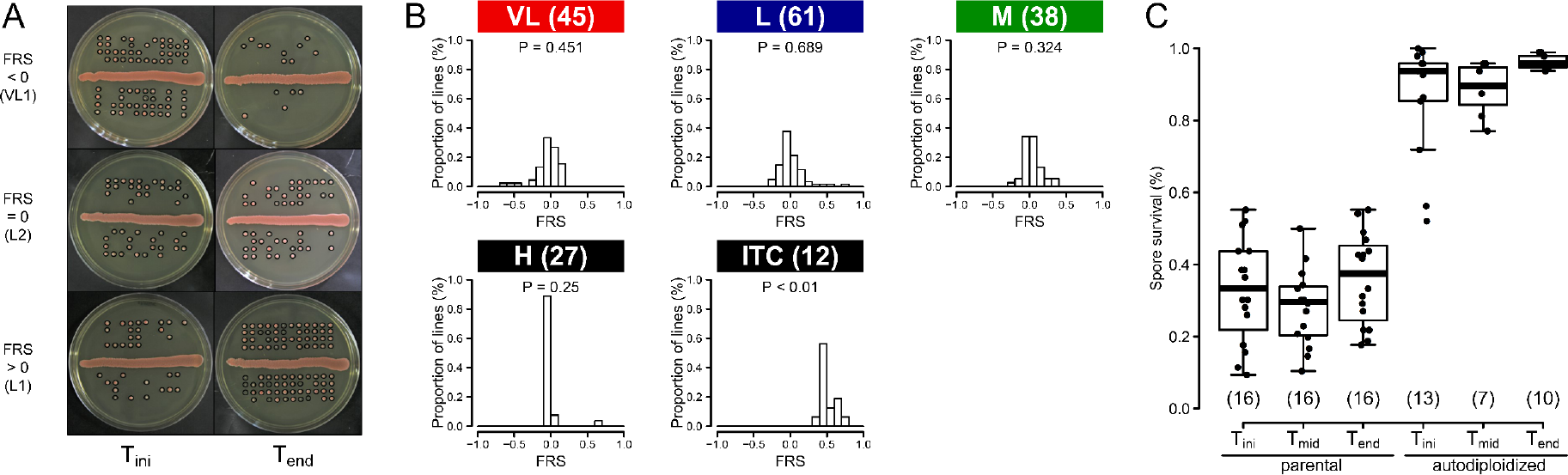
Hybrids gain and lose fertility at similar rates while sexual reproduction systematically improves fertility. (**A**) Example of spore survival from lines showing a reduction of fertility (top), no change of fertility (middle) and recovery of fertility (bottom). Spores considered as viable are circled in black. (**B**) Fertility recovery scores (FRS) show no clear directionality in mitotically propagated lines (VL_div_, L_div_, M_div_ and H_div_). P-values given for exact binomial tests. FRS for intra-tetrad crosses (ITC) indicate large and systematic increase in fertility. (**C**) Viable spores dissected from initial hybrids and evolved lines (parental) recover fertility to above 80% after autodiploidization. Numbers in parentheses represent sample sizes.

Although no general trend towards fertility recovery was observed during the experiment, we observed cases of spectacular fertility recovery in seven mitotic lines (4 L_div_ lines, 2 M_div_ lines and 1 H_div_ line), which displayed FRS close to that of ITC lines and similar to what is typically observed for diploid parents (*18*). For most of these lines, fertility restoration happened within 385 mitotic generations. As infertility of *Saccharomyces* hybrids is thought to be mainly due to recombination failures between homeologous chromosomes (*21*), there could be two main explanations for these sudden increases in fertility. Both explanations rely on extremely rapid loss of heterozygosity across the genome or potential correct chromosome pairing during meiosis. The first one is autopolyploidization, which would lead to spontaneous production of identical homologues that would facilitate chromosome pairing. The second is that the strains could have sporulated during the experiment and genetically compatible spores of opposite mating types could have mated. This event, although unlikely, would be the equivalent of autodiploidization. Mitotic loss of heterozygosity could also be involved but would not be expected to lead to such dramatic changes (*22*). We tested these hypotheses by measuring the total cellular DNA content of the lines to infer ploidy and by genome-wide genotyping.

We measured ploidy in all the lines at the three time points T_ini_, T_mid_ and T_end_ using flow cytometry. Surprisingly, these analyses revealed that some lines already deviate from diploidy after hybridization. While almost all the hybrids are diploid, both independent L_div_ hybrids show frequent triploidy (average at Tini of 54% triploid individuals per crosses) (Fig. 3A, B). It appears that this triploidization is a major driver of loss of fertility in the L_div_ hybrids (both crosses), with an average reduction of 45% compared to diploids (20.7% compared to 37.6% considering all time points, Fig. S8). We examined the genotype of hybrids using genotyping-by-sequencing (GBS) and found that at Tini, all triploid hybrids were composed of two copies of the *SpC* genome and one copy of the *SpB* genome (Fig. 3C). We also observed a small fraction of diploid cells in both parental *SpC* haploid stocks (Fig S9). These cells are pseudo-haploids, i.e. diploid but competent for mating (Fig S10). Triploidy appears to be frequent in the L_div_ and was observed in the two biological replicates performed with independent strains but not in all crosses between these two species (Fig. S11, S12) and is variable among replicates (Table S5). These results suggest that the stochastic duplication of the *SpC* genome happened prior to mating and that some *SpC* haploid strains used in this study may be prone to spontaneous genome doubling.

**Fig. 3.**
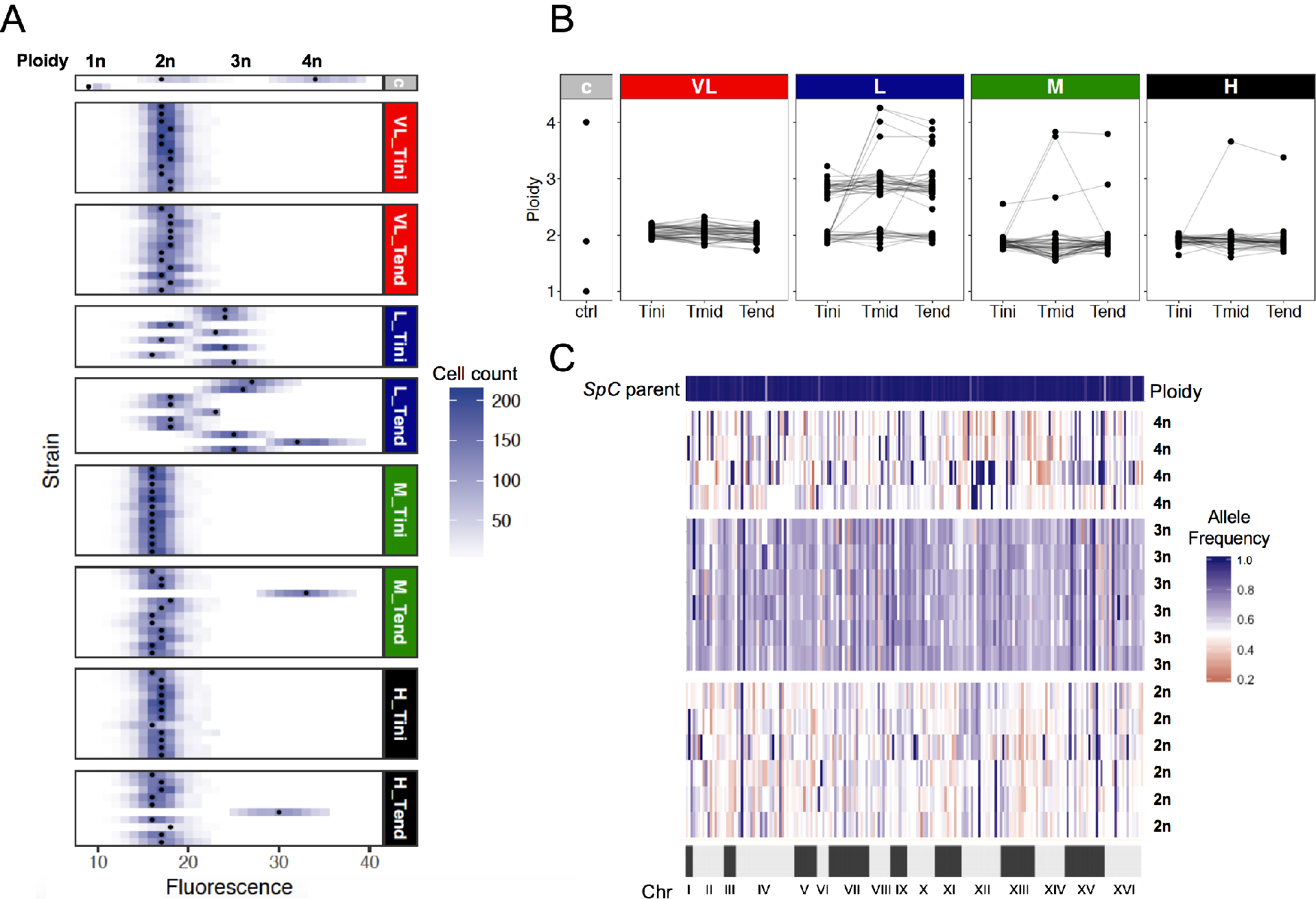
Ploidy varies among hybrids and evolves through time. (**A**) Ploidy of a subset of hybrid lines after hybridization (T_ini_) and after ~770 (T_end_) of mitotic generations. The c grey panel corresponds to controls. (**B**) Ploidy at the three tested timepoints (T_ini_, T_mid_ and T_end_). Connected dots represent independent lines. The c grey panel corresponds to controls. (**C**) Frequencies of 171 markers across the genome show around 50% of *SpC* alleles in the diploid and tetraploid strains and 66% in triploid strains.

Consistent with the hypothesis above based on polyploidy, the 6 lines showing more than 70% fertility after evolution became tetraploid (Fig. 4). This is also true even for hybrids between *S. paradoxus* and *S. cerevisiae*, which were initially completely sterile (Fig. 4). GBS analysis revealed that tetraploid hybrids result from a whole genome duplication of both parental genomes, since allele frequencies across 171 markers among the 16 chromosomes are around 50% (Fig. 3C, S13 and S14). It was shown before that artificially induced chromosome doubling in infertile hybrids between *S. paradoxus* and *S. cerevisiae* hybrids could restore fertility (*23*). Our results show that this happens spontaneously and without the need for natural selection.

**Fig. 4.**
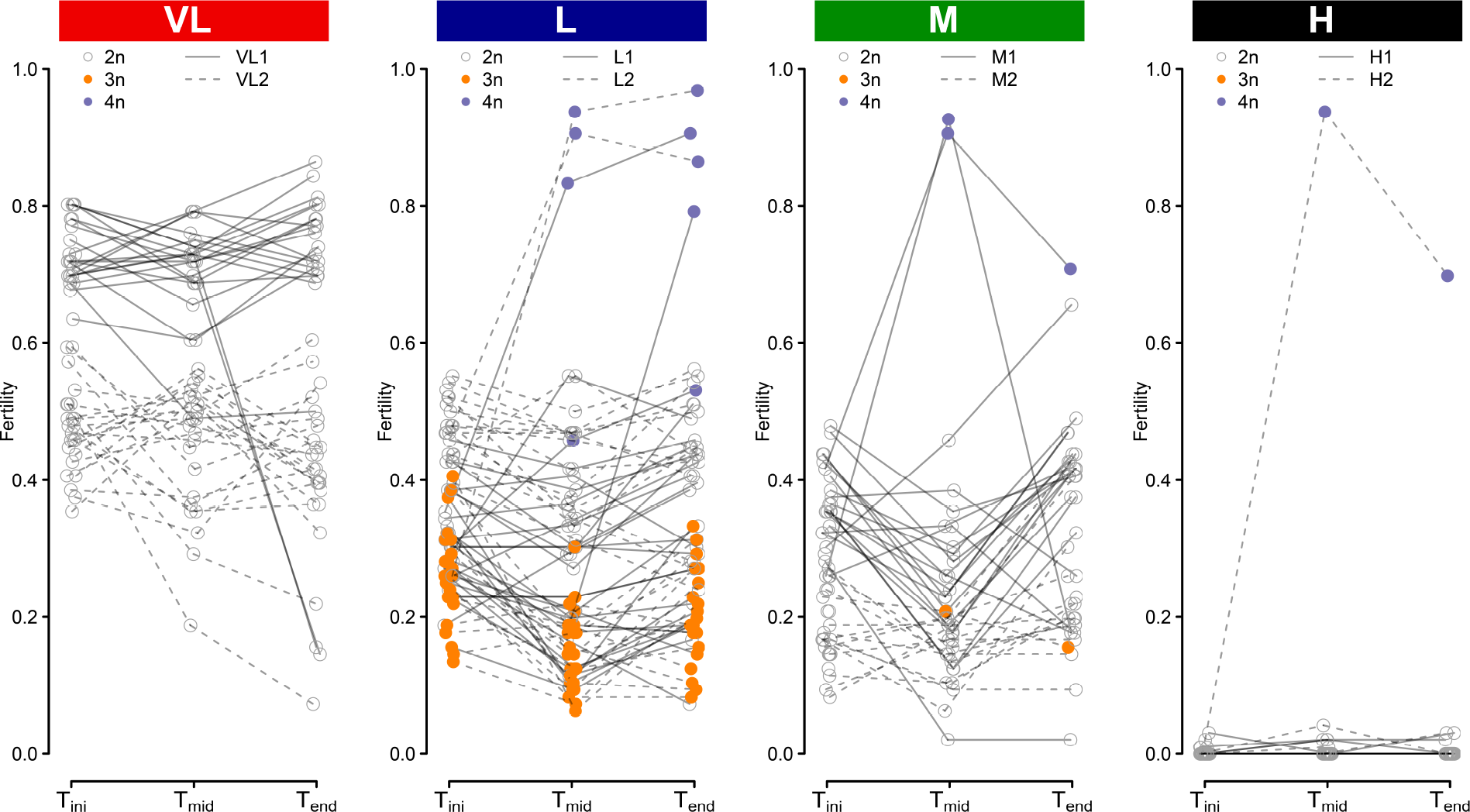
Triploidy leads to low initial fertility while tetraploidy leads to fertility recovery over time. Fertility trajectory at the three timepoints (T_ini_, T_mid_ and T_end_). Each connected dot represents an independent line. The colors correspond to ploidy. Dotted and full lines represent the two independent crosses within each genetic divergence class. In the case of VL1 and VL2, the two pairs of strains show different level of fertility, which is frequent in yeast intra-species crosses.

Our study shows that fertility can evolve in the absence of sexual reproduction, like a neutrally evolving quantitative trait, with incremental gains and losses, with no particular direction. However, in rare events, it can be recovered through changes of large effects. Mitotic fertility recovery contrasts with our results and that of others showing that sexual reproduction leads to incremental and rapid recovery of fertility (*22, 24*). However, sexual reproduction is thought to be extremely rare in yeast (*25*) and thus is unlikely to explain successful hybridization and introgression in the wild. Our data shows that hybrids can recover fertility by becoming allopolyploids in less than 400 cell divisions, which could represent less than 250 days in nature (*18*). Fertility recovery could therefore happen before sexual reproduction occurs. How frequently this occurs in nature remains to be tested. However, it could contribute to important events. For instance, it was shown recently that the whole-genome duplication (*26*) in an ancestor of the *Saccharomyces* genus originated from inter-species hybridization event. The success of such an event would have been unlikely if the F1 hybrids were completely sterile and if sexual reproduction would have been needed to recover fertility. Our results suggest that whole-genome duplication could have happened spontaneously and neutrally, thus restoring fertility at the same time. We also find triploidy in some of our crosses, which greatly reduces hybrid fertility. Ploidy changes are therefore a double-edged sword, causing both reproductive isolation and fertility recovery.

## Supporting information

DataS1

## Acknowledgments

We thank the members of the Landry lab for discussions and J. Hallin, A. K. Dubé, C. Eberlein, A. Fijarczyk, N. Aubin-Horth, J.B. Anserson, L. Kohn, S. Otto and A.M. Dion-Coté for comments on the manuscript.

## Funding

NSERC Discovery grant and Canada Research Chair to C.R.L, FRQNT scholarship to G.C and M.H, NSERC Alexander Graham-Bell scholarship to G.C. and FRQS post-doctoral fellowship to S.M;

## Author contributions

Conceptualization, C.R.L., G.C., M.H., S.M.; Data curation, G.C, S.M and M.H; Funding acquisition, C.R.L.; Experimental work and formal analysis: Cell propagation by G.C., S.M., M.H., Fertility by G.C., Ploidy and GBS by S.M., Mitochondrial DNA by M.H, R scripts support for ploidy/GBS by H.M. Project supervision, C.R.L.; Writing - original draft, G.C. and S.M.; Writing – review & editing, all authors;

## Competing interests

Authors declare no competing interests.

## Data and materials availability

Raw sequencing data can be accessed under BioProject Accession # PRJNA515073. All yeast strains are available from C.R.L. under a material transfer agreement.

## Materials and Methods

### Strain construction

To ensure that the evolved strains would keep their mitochondrial genome through the experiment, we used the *ade2*-*Δ* marker to help with visual identification of respiration deficient colonies, a strategy used in past mutation accumulation experiments (*27, 28*). The heterothallic *S. paradoxus* strains were generated previously (*18*, *29*, Table S1). The *ADE2* and *HO* loci of the two wild *S. cerevisiae* strains were deleted following the method described by Güldener *et al*. (1996)(*30*). The *ADE2* locus of the *S. paradoxus* strains were replaced by homologous recombination with resistance cassette following the same procedure as for *HO* in *S. paradoxus* (*29*). Oligonucleotides with overhangs (Table S3) specific to each lineage were used to generate the deletion cassettes from pFA-hphNT1 (*31*) to prevent recombination with the cassettes already present at the *HO* locus (*KANMX* and *NATMX* cassettes).

### Experimental crosses

Two crosses were made for each of the divergence levels (L1, M1, H1, VL1 and L2, M2, H2, VL2) (Table S3). All incubation steps were performed at room temperature (RT). Haploids to be crossed were precultured overnight in 5 mL of YPD (1% yeast extract, 2% tryptone and 2% D-glucose). Precultures were then diluted at OD_600nm_ of 1.0 in 500μL aliquots. The aliquots from two strains to be crossed were mixed together and 5μL were used to inoculate 200μL of fresh YPD medium in 96 replicates so all strains would derive from independent mating events and would be truly independent hybrids. Cells were given 6 hours to mate after which 5μL of the mating cultures were spotted on a diploid selection medium (YPD, 100μg/mL G418, 10μg/mL Nourseothricin). From each of the 96 spots, 1 colony was picked as a founding line for the evolution experiment, resulting in 96 independent lines for each of the six interlineage crosses (48 lines for the two intra lineage crosses).

### Evolution experiment

Each of the independent lines (single colonies) were streaked on one third of a YPD agar plate. To facilitate the detection of potential lines mixing during the experiment, each Petri was streaked with three different crosses (series L1, M1, H1 and L2, M2, H2). Crosses VL1 and VL2 were streaked on two different sets of Petri dishes, with three lineages per Petri. The 192 plates were split into 3 sets of 64 (lines 1-64 and lineages 65-96). Plates were incubated at room temperature for 3 days after which a new single colony was streaked as a progenitor for the new generation. Each set was rotated between three manipulators at each replication steps. The criteria for the new colony were to be 1) the closest to a predesigned mark on the Petri dish, allowing for unbiased colony selection, 2) a single colony and 3) big enough to allow for both replication on a new medium and the inoculation of a liquid culture to generate a frozen stock. If the colony closest to the mark did not meet criteria 2 and 3, the second closest colony was then examined, and the process was repeated until all criteria were met. Every 3 passages, the colonies were both streaked and used to inoculate the wells of a 96 wells plate containing 150μL of fresh YPD medium. After a 24 h incubation at room temperature, 75 μL of 80% glycerol was added and the plates were placed in a −80°C freezer for archiving. The lines were maintained on plates for a total 35 passages.

### Estimation of generation time

To evaluate the generation time on plates and thus estimate the total number of mitotic divisions during the experiment, 3 lines from each cross were randomly selected. Strains for T_ini_ and T_end_ were thawed (48 total strains tested), streaked on YPD solid medium and let grow for 3 days. Strains were then replicated on fresh YPD solid medium following the same protocol as for the evolution experiment. After 3 days of incubation, the colony closest to the predesigned mark was extracted from the media using a sterile scalpel. The agar block with the colony was put in a sterile 1 mL Eppendorf tube and the colony was resuspended in 500μL of sterile water. Optical density at 600nm (OD_600nm_) of the resuspensions was estimated using a TECAN Infinite 200 plate reader (TECAN, Männedorf, Switzerland). These resuspensions were diluted in 200μL of sterile water to obtain OD_600nm_ values of about 0.05 (500 cells/μL). The dilutions were then analyzed with a Guava^®^ easyCyte HT (Millipore Sigma, Burlington, USA) flow cytometer to estimate actual cell numbers. The estimated number of cells/μL were used to calculate the initial number of cells in the volume used in the dilution and then in the initial 500μL. The log_2_ of this number represents the number of cell doubling during the colony growth for 1 passage of the experiment assuming the colony was formed from a single cell (Fig. S5, Data S1).

### Sporulation protocol

Strains were thawed and 2μL of the stocks were spotted on a fresh YPD medium and incubated for 3 days. A small number of cells was used to inoculate 4 mL of fresh YPD media and incubated for another day. From those precultures, a new 4 mL culture was inoculated at 0.6 OD_600nm_ in fresh YPD and grown for 3 hours. Cultures were then centrifuged at 250 g and the YPD was replaced with 4mL of YEPA medium (1% yeast extract, 2% tryptone and 2% potassium acetate). Cultures were incubated for 24 h after which they were centrifuged again at 250 g, washed once with sterile deionized water and put into 4 mL of SP medium (0.3% potassium acetate 0.02% _D_-Raffinose). After 3-5 days of incubation, the strains were dissected as in Charron et al. 2014 with a SporePlay™ dissection microscope (Singer Instruments, Somerset, UK) on YPD plates and incubated for 5 days. Pictures of the plates were taken after the incubation and fertility was determined as the number of spores forming a colony visible to the naked eye after 5 days.

### Mitochondrial DNA genotyping

Two mitochondrial loci were genotyped for presence or absence by PCR. Total DNA extractions were performed using the method described by Looke *et al*. 2011 (*32*). The two PCR assays target loci in the *RNL* and *ATP6* mitochondrial genes, respectively, as described in Leducq *et al*. 2017 (*33*). Multiplex PCR with both primer pairs (Table S2) was performed with the following cycle: 3 min at 94°C; 40 times the following cycle: 30 s at 94°C, 30 s at 57.5°C, and 50 s at 72°C; and 10 min at 72°C. A PCR targeting the ITS1-5.8S-ITS2 locus was performed on the same DNA samples as positive controls following the method described in Montrocher *et al*. 1998 (*34*).

### Fertility assessment in the evolved lines

Fertility of the evolved lineages (VL, M and H) was measured on 24 randomly chosen lines per cross (14 which lived through P35 and 10 that were lost before the end of the experiment). To ensure we had the same numbers of diploids to compare with the other lines, more strains were chosen for the two L crosses, 36 lines were used (24 diploids and 12 triploids, 26 strains that lived until P35 and 10 lost before). For each line, we measured fertility at three different time points: 1) immediately after mating (T_ini_), 2) at the halfway point for the given lineage (T_mid_) and 3) at the last passage of the given line (T_end_). This means that Tmid and Tend do not always refer to 385 and 770 mitotic generations. Fertility data is available in the Data S2 file.

### Autodiploidization of spores

In order to generate fully homozygous strains that should have fully recoverd fertility, we performed autodiploidization experiments. After the dissections, some spores from the L and M crosses were typed for their mating type locus and their antibiotic resistance markers. When possible, four spores were kept as frozen stocks (two of each mating type and resistance combination). A subset of the spores expressing the G418 resistance was selected to undergo autodiploidization. This was performed by transformation of the spores with the plasmid pHS3 containing *S. cerevisiae HO* gene with its endogenous promoter and the CloNAT resistance cassette (pHS3 was a gift from John McCusker, Addgene plasmid # 81038). Transformants were selected on fresh selection medium (YPD, 200μg/mL G418, 100μg/mL CloNAT) to be sporulated and dissected.

### Intra-tetrad crosses (ITC)

Intra-tetrad mating was conducted in order to generate hybrids with rapid loss of heterozygosity and to test whether fertility recovery was possible. Strains from the L1 and the L2 crosses were sporulated as described above, but the dissection steps were modified to allow the mating of pairs of spores from the same tetrad (leaving 2 spores instead of one at the designed dissection spot). The plates were incubated for 5 days. In this manner, pairs of spores of opposite mating types could mate to generate new diploids while pairs of identical mating types would divide mitotically as haploids. To ensure that the yeasts recovered were the result of a mating event, we selected diploids with resistance to both G418 and CloNAT by replica plating the colonies on a fresh selective medium (YPD, 200μg/mL G418, 100μg/mL CloNAT). From the surviving diploids, one colony was randomly selected to be sporulated again. This process was repeated 12 times (ITC 1 to ITC 12) for 16 lines (2 replicates from lines L1-6, L2-8, L2-36 and L2-63 and 8 other unique line). This number of meiosis was expected to generate extensive LOH as, for a single given heterozygous locus, less than 1% of the population will have maintained heterozygosity (*35*). The ITC 0 (initial hybrids), ITC 1, ITC 6 and ITC 12 strains were sporulated and dissected as described above.

### Determination of ploidy

Measurement the cell DNA content was performed using flow cytometry with the SYTOX™ green staining assay (Thermofischer, Waltham, USA). Cells were first thawed from glycerol stocks on solid YPD in omnitray plates (room temperature, 3 days) including controls. The parental strain *SpB* (MSH604) was used as control on both its haploid and diploid (wild strain) state. Liquid YPD cultures of 1 ml in 96 deepwell (2ml) plates were inoculated and incubated for 24h at room temperature. Cells were subsequently prepared as in Gerstein et al (2006)(*36*) but stained with a final SYTOX™ green concentration of 0.6 μM for a minimum of 1 h at room temperature in the dark. The volume of cells was adjusted to be around a cell concentration less than 500 cells/μL. Five thousand cells for each sample were analysed on a Guava^^®^^ easyCyte 8HT flow cytometer using a sample tray for 96-well microplate. Cells were excited with the blue laser at 488 nm and fluorescence was collected with a green fluorescence detection channel (peak at 512 nm). The distributions of the green fluorescence values were processed to find the two main density peaks, which correspond to the two cell populations, respectively in G1 and G2 phases. The data was analysed using R version 3.4.1 (*37*).

### Genotyping by sequencing

We performed genotyping-by-sequencing (GBS) to investigate the genomic composition of the triploid and the tetraploid hybrids. We sampled 77 strains in total: 8 diploids and 16 triploids from (L) crosses at T_ini_; all 8 tetraploids and as controls 2 diploids from each cross L_div_, M_div_ and H_div_ at T_ini_ and T_end_ as well as all parental strains. DNA was extracted from overnight cultures issued from one isolated colony following standard protocols (QIAGEN DNAeasy, Hilden, Germany). As controls, we prepared artificial hybrid genomes by mixing DNA of parental strains with different proportion from each 0.5/0.5, 0.66/0.33 or 0.33/0.66. DNA was quantified using Accuclear^^®^^ Ultrahigh sensitivity dsDNA Quantification kit (Biotium, Fremont, USA) in a Spark^^®^^ microplate reader (TECAN, Männedorf, Switzerland). DNA concentration was normalized to 10 ng/μl and subsequently used for library preparation.

Libraries for Ion Proton GbS were prepared using the procedure described by Masher et al., 2013 (*38*) at the Plateforme d’Analyses Génomiques of the Institut de Biologie Intégrative et des Systèmes (IBIS, Université Laval, Québec, Canada) with the following modifications: ApeKI endonuclease enzyme and ApeKI barcodes were used instead of the PstI/MspI combination and a blue Pippin (SAGE science, Beverly, USA) was used to size libraries (150- 300bp) before PCR amplification. Libraries were prepared for sequencing using an Ion Chef™, Hi-Q reagents and PI™ chip kit V3 (ThermoFisher, Waltham, USA) and the sequencing was performed for 300 flows. A single fastq file was obtained and demultiplexed using Radtags tool from STACKS with default options (*39*) which generated separated fastq files for each sample. Reads were mapped onto the *S. paradoxus* reference genome (CBS432) (*40*) for all crosses and onto the *S. cerevisiae* reference genome (YPS 128) (*40*) for H cross using Bowtie2 (*41*). Read coverage for each position in the genome was also determined using SAMtools depth. Single nucleotide polymorphism (SNP) calling on these GBS libraries was performed using the BCFtools from SAMtools (*42*) with default parameters. The data was subsequently analysed using R version 3.4.1 (*37*). To measure the allelic frequency, we filtered for SNPs with more than 10X coverage and corresponding to positions in the genome covered in both parental strains to be able to identify the parental origin of the alleles in hybrids.

### Statistical analyses

Survival curves were produced and analyzed using the R packages *survival (43)* and *survminer* (*44*). The analysis of the correlation between sporulation and genetic divergence was performed using the glm R function to perform a logistic regression with the formula: Sporulation T_end_ ~ Genetic Divergence. Statistical analyses and figure creation for fertility data were done using custom scripts (Data S3) in R version 3.3.2 (*37*). Figures and statistical analyses for mtDNA loss and sporulation capacity were performed using custom scripts in Python (version 3.6.3).

## Supplementary Text

### *ade2-Δ* colony coloration phenotype

Although we used the *ade2*-*Δ* marker as a visual aid to track loss of mitochondrial DNA, we still passaged strains that seem to have loss mitochondrial DNA. During the experiment, some colonies from all crosses suddenly turned whitish or light orange. This paler coloration did correlate with the absence of sporulation in the strains. The only thing that changed during the evolution experiment is the yeast extract (EMD millipore, Burlington, USA), for which the lot number changed. Further testing suggests that the pink/red coloration of the *ade2*-*Δ* mutants is media dependent. On one of the yeast extract lot used, white colonies appear red and show slower growth while on the other, most colonies are white and show normal growth. The slower growth is common to all *ade2*-*Δ* strains (Fig. S4A, B).

**Fig. S1.**
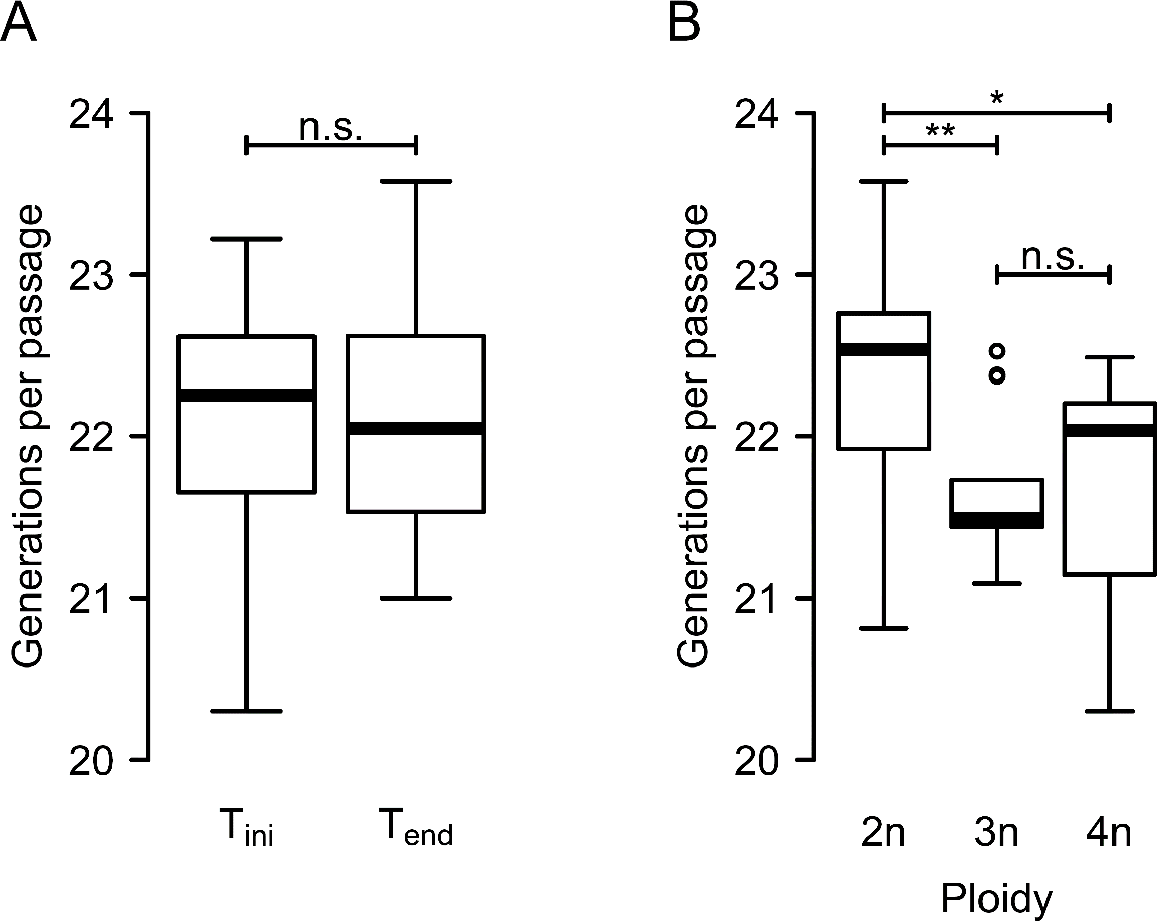
Number of mitotic generations per passage is stable through time but differ between hybrids with different ploidies. Calculated number mitotic generations as the log_2_ of the number of cells in colonies obtained by following the experimental procedure of the evolution experiment for a subset of 40 lines. (**A**) There is no statistically significant difference in the number of mitotic mitotic generations when considering the timepoints (T_ini_ or T_end_) of the experiment (one-way ANOVA (F(1,61) = 0.015, P = 0.902)). (B) but there is a significant effect of ploidy of the hybrids lines (Tukey HSD, “**” P < 0.01, “*” P < 0.05, “n.s.” non-significant) (**B**).

**Fig. S2.**
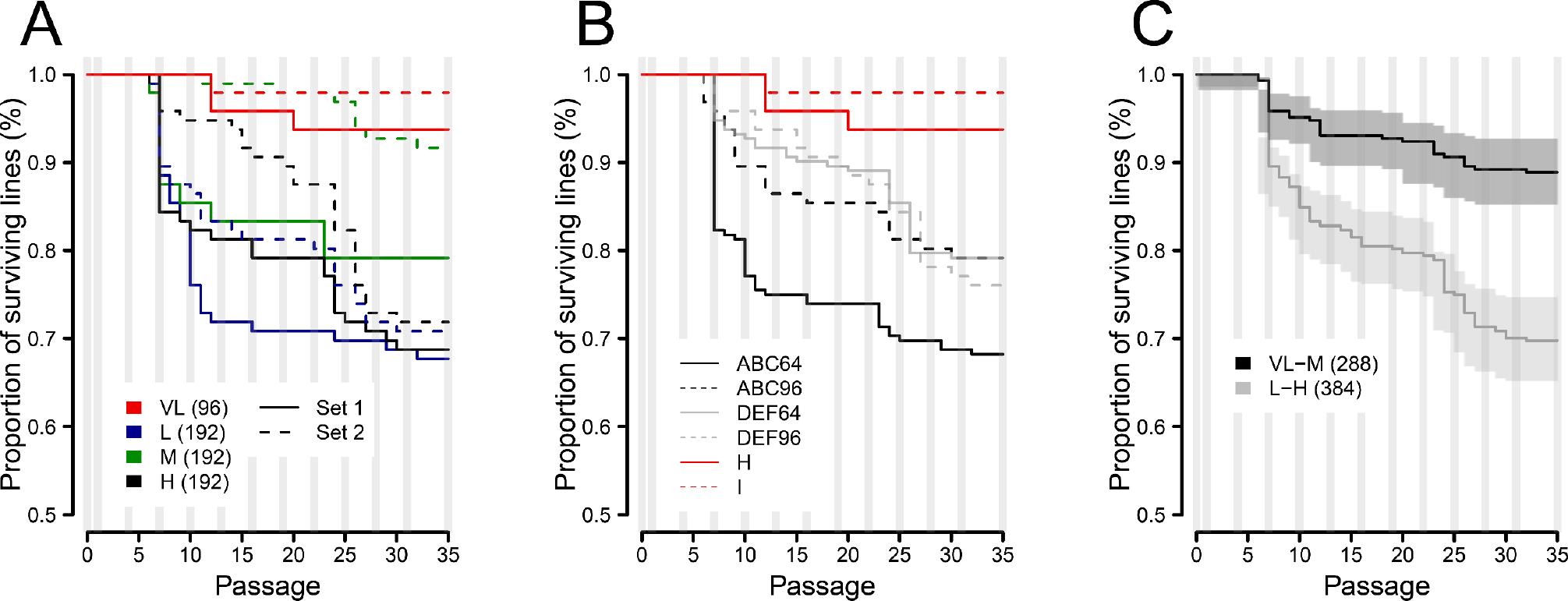
Survival of the evolution lines. Survival of the lines grouped by (**A**) individual crosses, (**B**) replication sets used during the evolution experiment and (**C**) crosses showing high (VL-M) and low (L-H) survival.

**Fig. S3.**
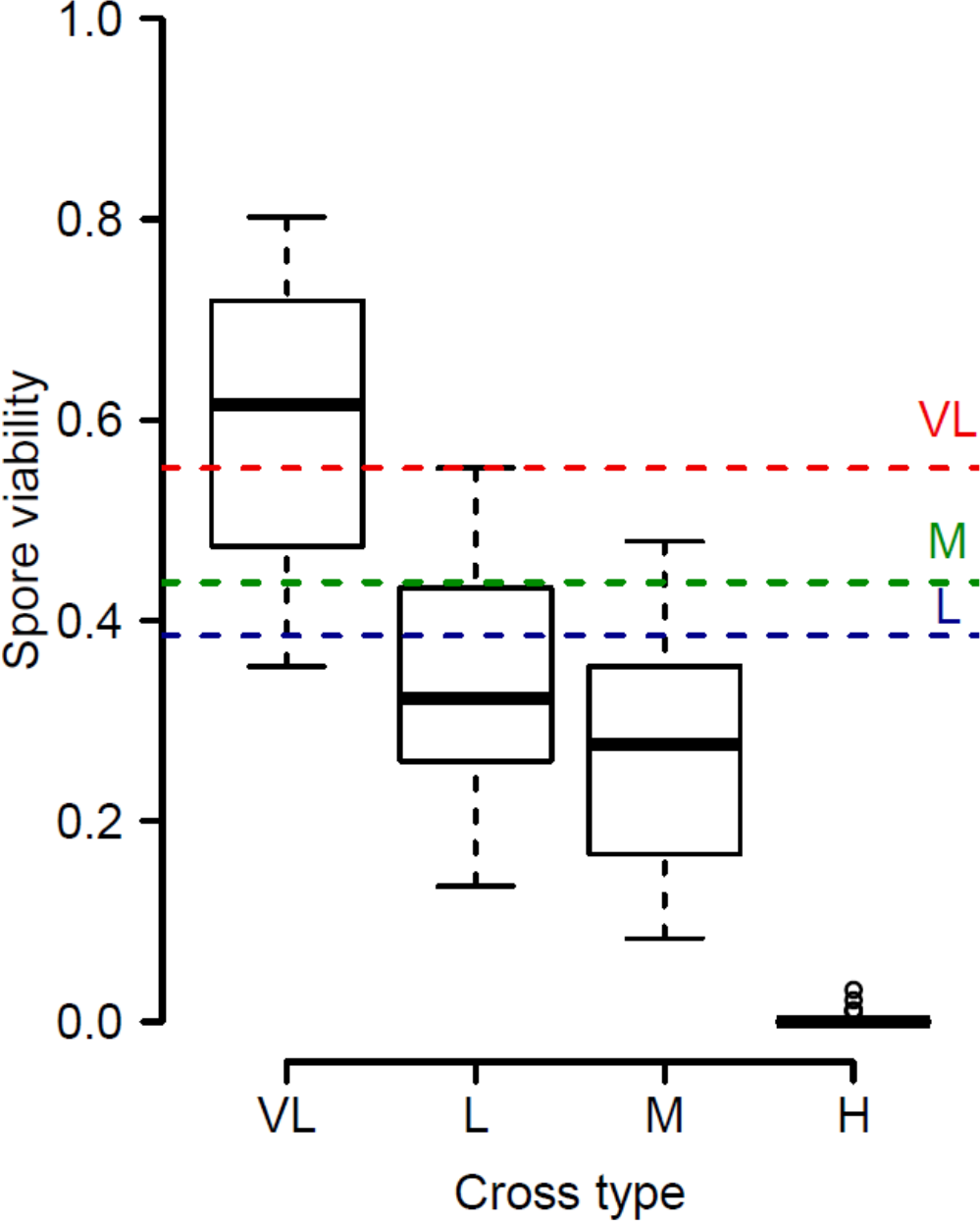
Ancestral lines show expected spore viabilities for their cross types. Boxplot of spore viabilities for the different hybrid lines at T_ini_. Dashed lines represent median spore viabilities values for each of the cross types from Leducq *et al*. 2016 (*18*).

**Fig. S4.**
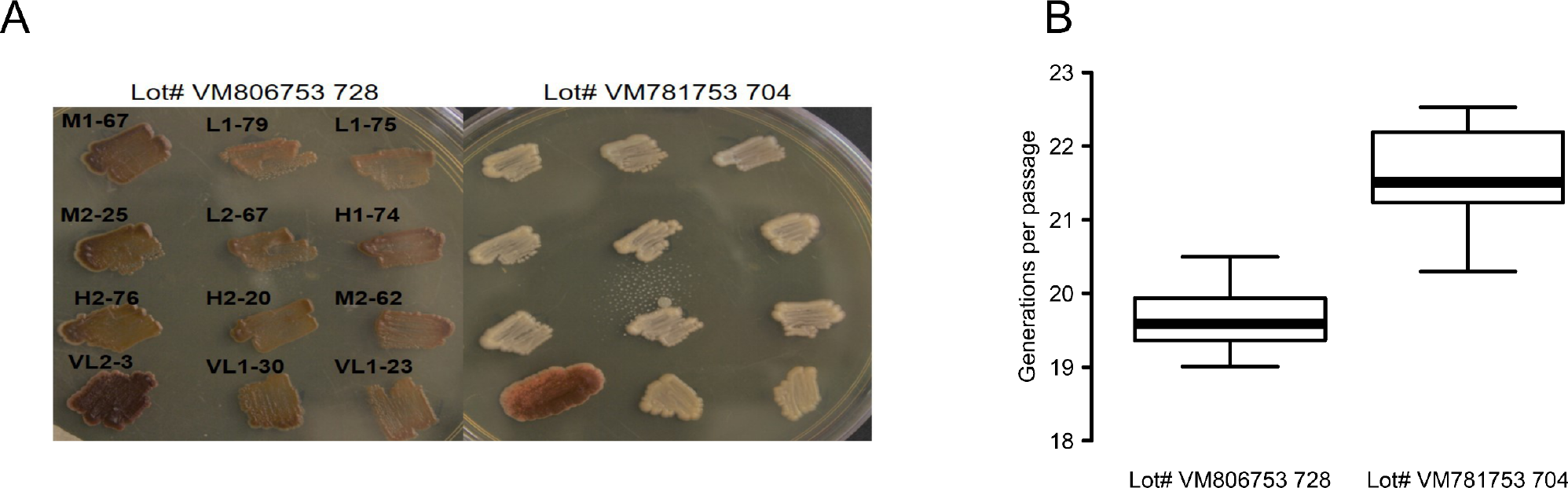
The *ade2*-*Δ* coloration phenotype and growth rate is media dependent. (**A**) Sporulation negative strains appear to be red when grown on YPD prepared with yeast extract from one lot (panel A left). Using the same ingredients but changing the lot number of the yeast extract yielded a very different coloration for all tested lines but one (panel A right). Evolution line numbers indicated above the cell patches. Plates were prepared and inoculated on the same day. Photos were taken after 5 days of growth. (**B**) Generations per passage on the two different media measured by cytometry on a subset of triploids and tetraploids that were sporulation positive.

**Fig. S5.**
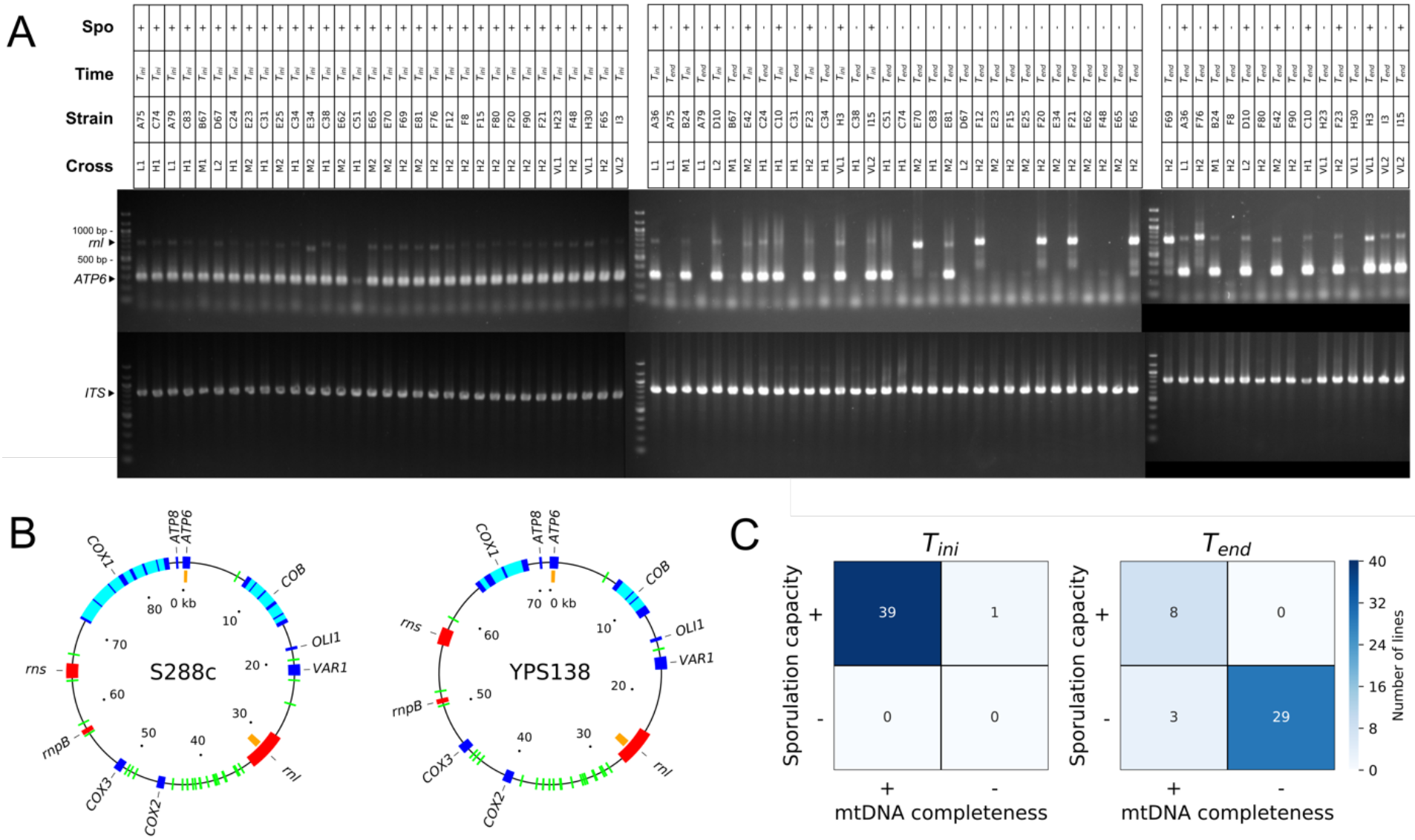
Partial or complete mtDNA deletion is associated with loss of sporulation capacity after 770 mitotic generations. (**A**) PCR assays for the *rnl* and *ATP6* mitochondrial loci and for the *ITS* nuclear locus. Presence or absence of these loci was assayed for 32 lines which lost sporulation capacity at the end of the mitotic evolution experiment and 8 lines that maintained sporulation capacity. PCRs were performed on DNA extracted from stocks from T_ini_ and T_end_ timepoints. The *rnl* amplicon is expected to vary in size around 700 bp, while the *ATP6* amplicon is expected to be 285 bp long. (**B**) Map of *S. cerevisiae* (S288c) and *S. paradoxus* (YPS138) mtDNAs. The rnl and ATP6 amplicons are shown in orange. Protein-coding genes are shown in blue, protein-coding gene introns in cyan, RNA-coding genes in red and tRNAs in green. The genome annotations used are from Yue et al. 2017 (*40*). (**C**) Partial or complete loss of mtDNA is associated with loss of sporulation ability. For T_ini_ and T_end_, contingency tables show the counts of lines according to mtDNA completeness (+: both markers present, -: at least one marker absent) versus sporulation capacity. The association is significant for T_end_ (Fisher’s exact test, odds ratio > 77, P-value=2.15×10^−6^).

**Fig. S6.**
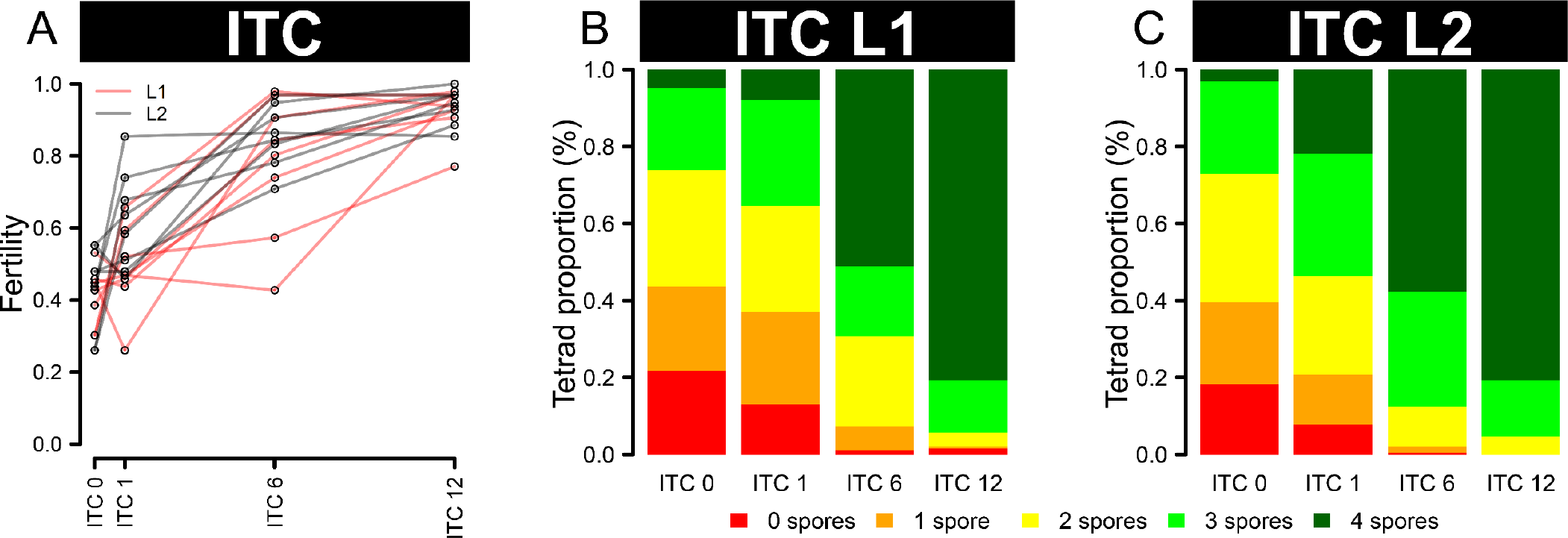
Intra-tetrad mating allows rapid restoration of hybrid fertility. (**A**) Fertility trajectories of 16 hybrids through 12 sporulations followed by intra-tetrad mating events. Line colors indicate the SpB×SpC cross identity. (**B**) and (**C**) Combined proportions of each tetrad types per cross type right after hybridization (ITC 0) and after 1, 6 and 12 intra-tetrad cross rounds.

**Fig. S7.**
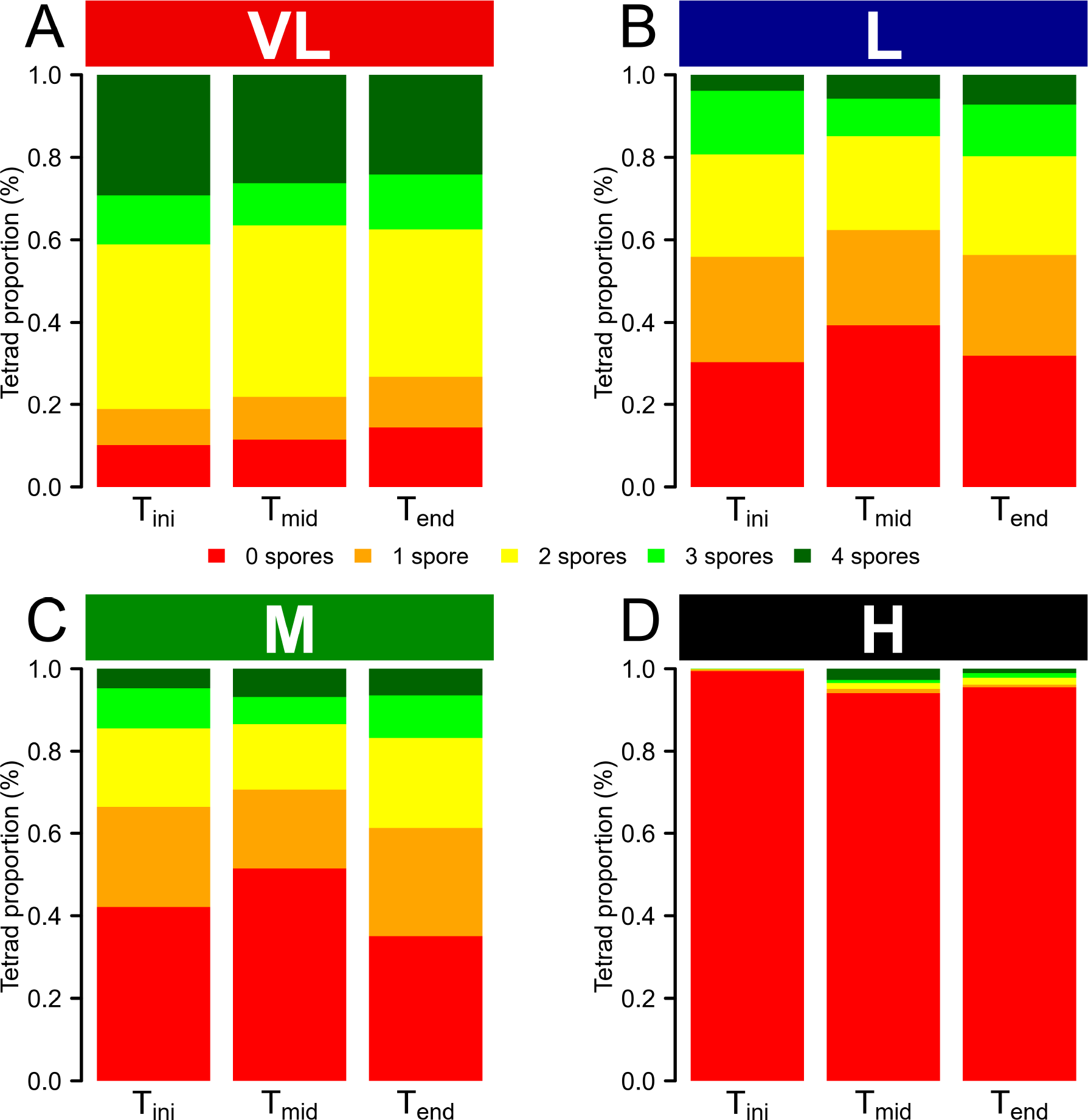
Tetrad type proportions do not change during evolution. The proportions of possible tetrad types at each time point tested for the (**A**) VL_div_, (**B**) L_div_, (**C**) M_div_ and (**D**) H_div_ crosses. The data of the two independent crosses for each cross type were merged for this analysis.

**Fig. S8.**
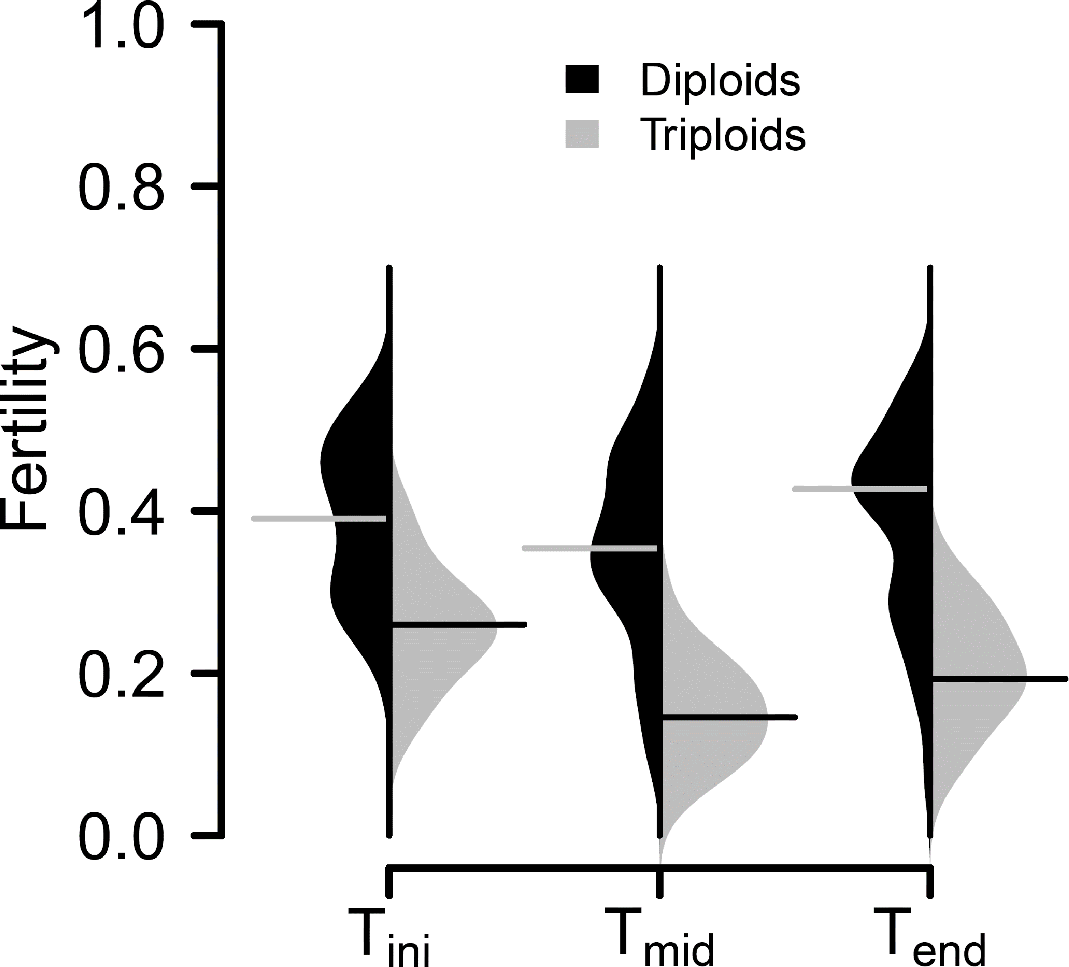
Triploid hybrids have a lowered fertility. Fertility value distributions for the diploids (black) and triploid (grey) individuals from the L lines at each of the three tested timepoints.

**Fig. S9.**
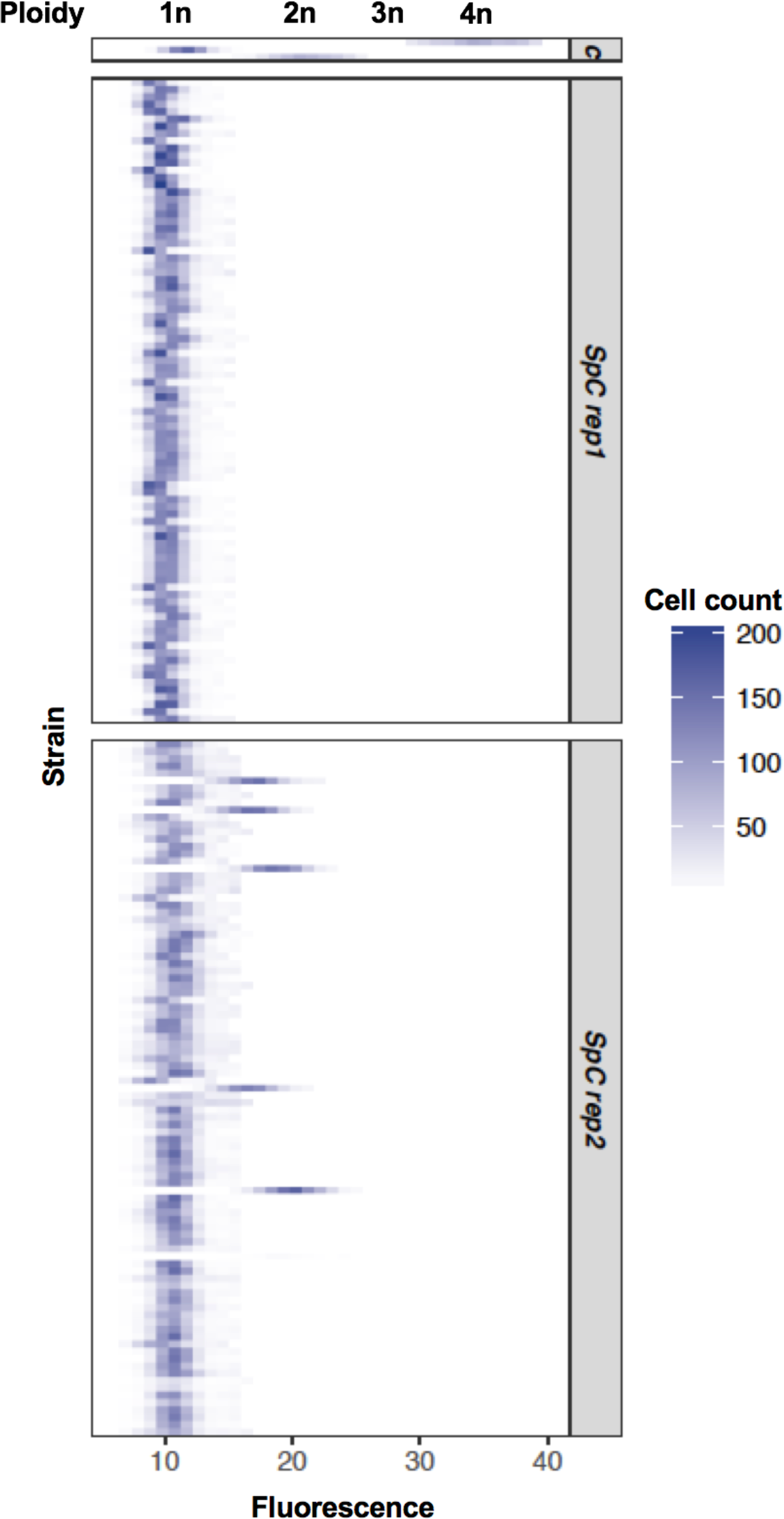
The parental *SpC* haploid stocks contain a small fraction of diploids. Ploidy of 94 isolated colonies from the parental *SpC* haploid stocks using flow cytometry repeated at two independent times. The c panel indicates controls.

**Fig. S10.**
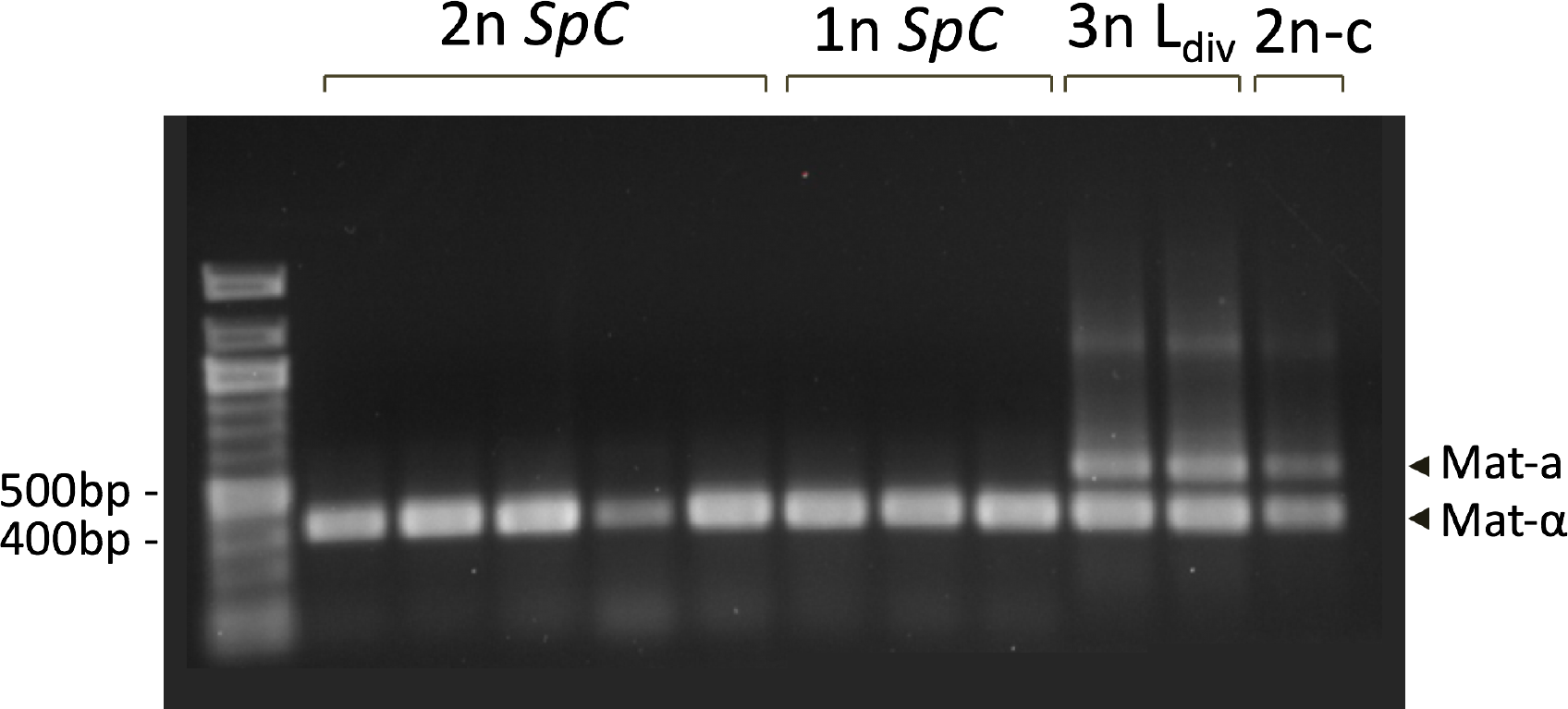
The small fraction of diploids in the *SpC* haploid stocks are pseudo-haploids. PCR of the mating type locus made on genomic the DNA of isolated colonies. The 5 diploids (2n *SpC*) identified in the *SpC* haploid stocks are shown along with 3 haploid *SpC* (1n *SpC*) from the same stocks, two verified triploid L_div_ lines (3nL_div_) and a diploid control (2n-c).

**Fig. S11.**
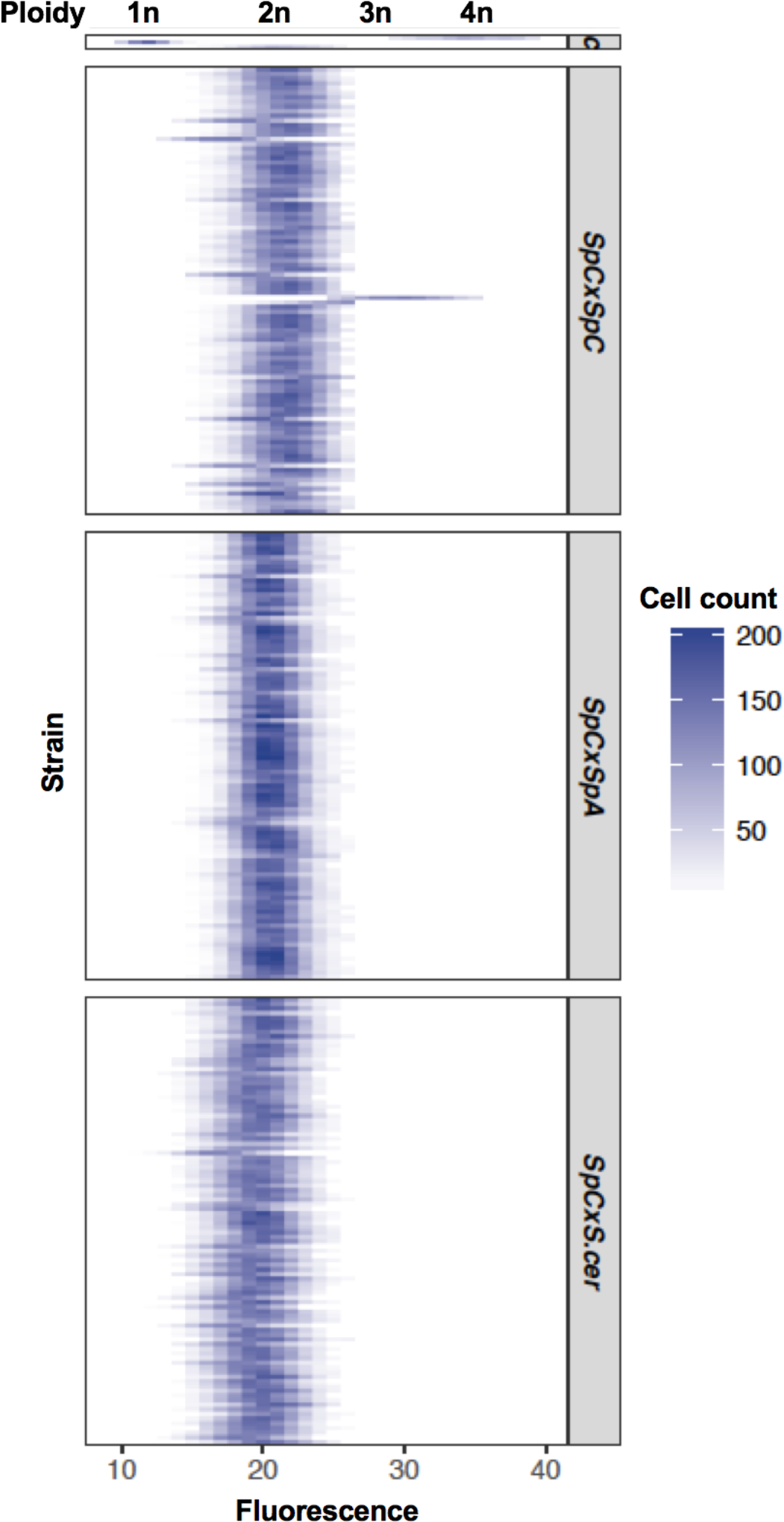
Triploidy is absent or rare from hybrids from crosses between *SpC* and other lineages and species. Ploidy variation after the hybridization of 94 independent replicates of (*SpC* x *SpC*), (*SpC* x *SpA*) and (*SpC* x *S. cerevisiae*) crosses using flow cytometry. The c panel corresponds to controls.

**Fig. S12.**
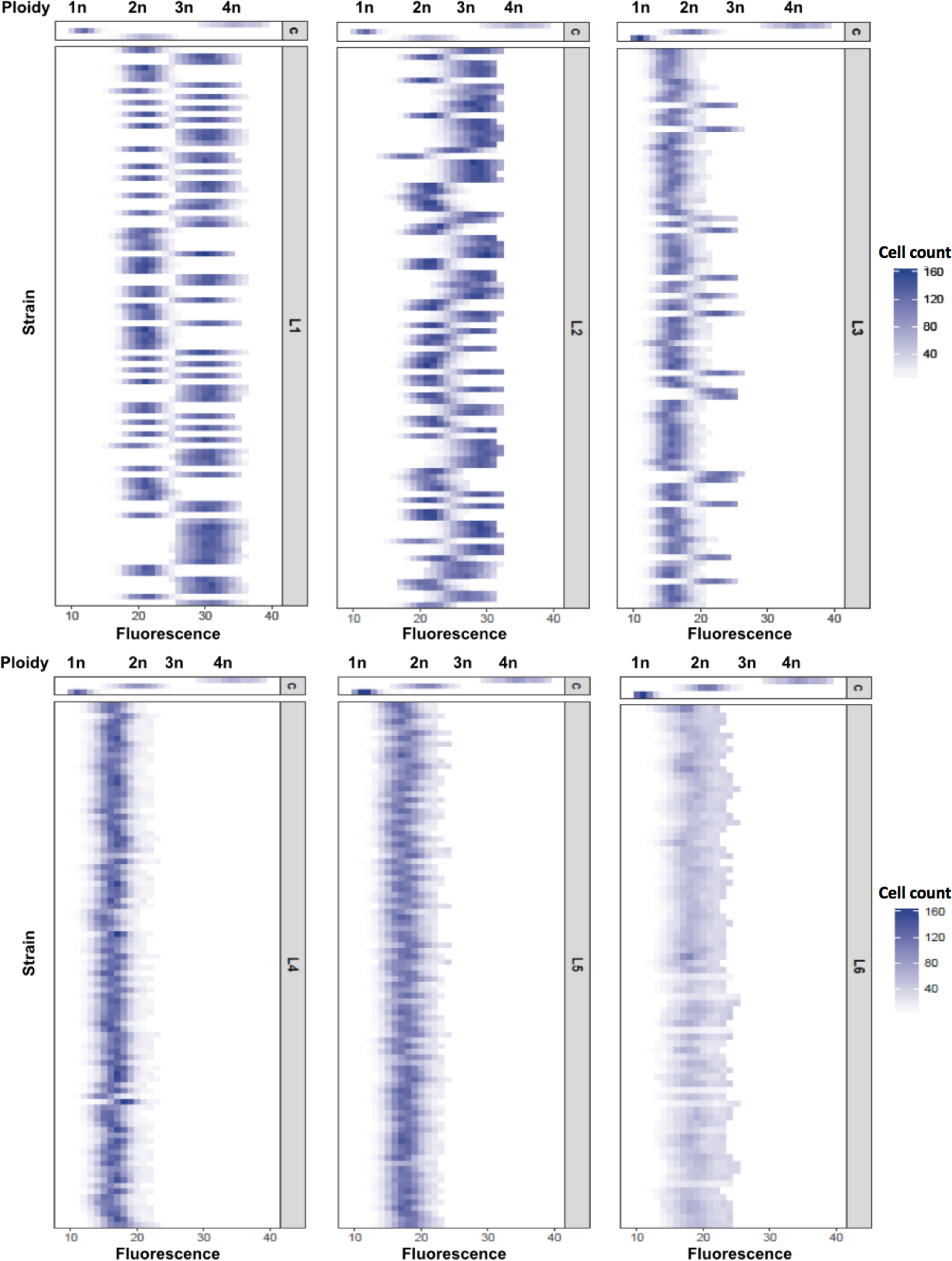
Triploidy is observed in crosses between different *SpB* and *SpC* strains but not in all crosses, revealing a stochastic origin. Ploidy variation after the hybridization of 94 independent replicates of different *SpB×SpC* crosses (L_div_1, L_div_2, L_div_3, L_div_4, L_div_5 and L_div_6) using flow cytometry. The c panel corresponds to controls. The strains used for each cross are listed in the Table S1 and S3.

**Fig. S13.**
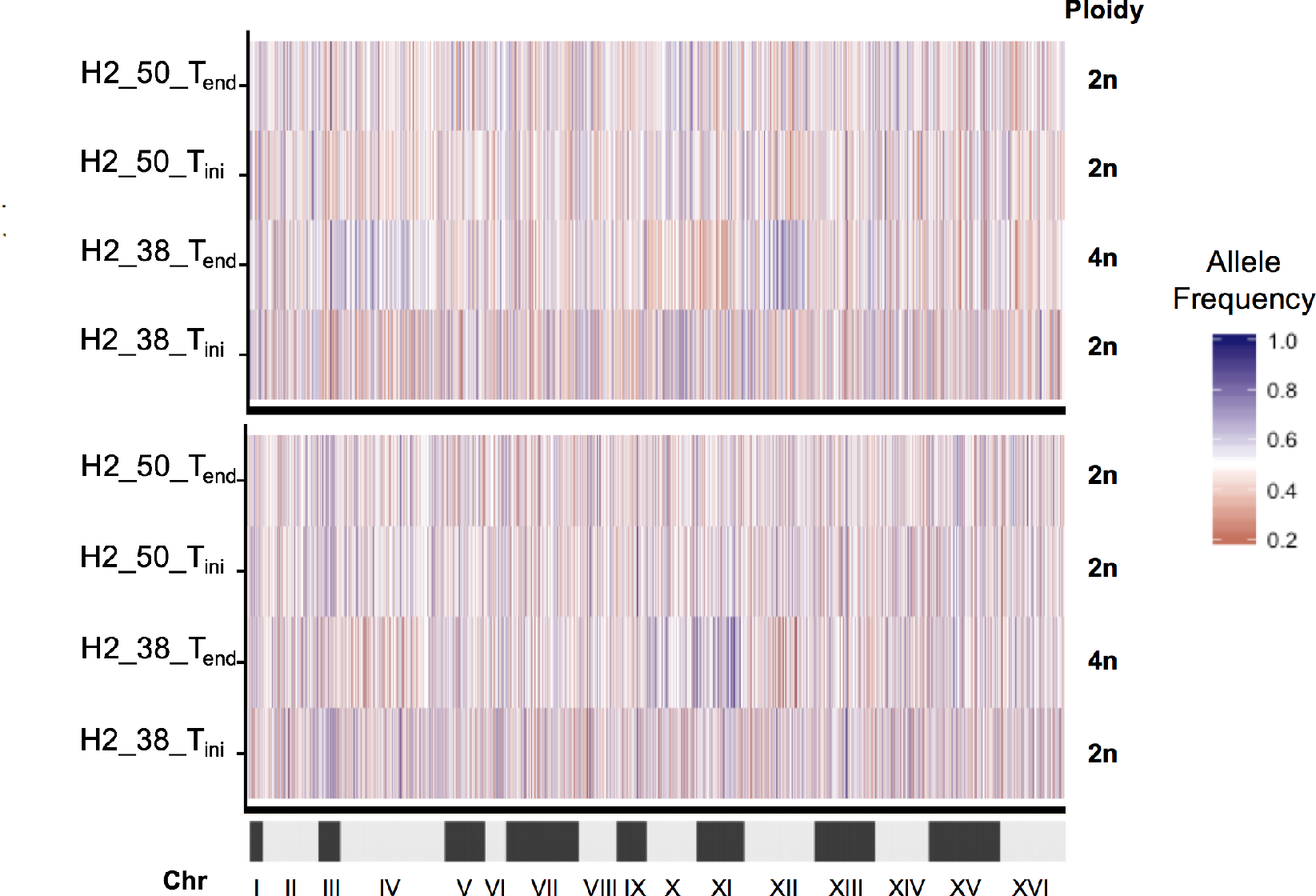
The tetraploid hybrid between *S. paradoxus* and *S. cerevisiae* result from a whole genome duplication of both parental genomes. Allele frequencies among the 16 chromosomes of the tetraploid H2_38 at T_end_ are around 50%, similar to what is seen for the H2_38 at T_ini_ and the diploid hybrid H2_50 at T_ini_ and T_end_. The heatmap on the top represent the allele frequency after mapping on *S. cerevisiae* reference genome (53078 markers) and the one on the bottom after mapping on *S. paradoxus* reference genome (56280 markers).

**Fig. S14.**
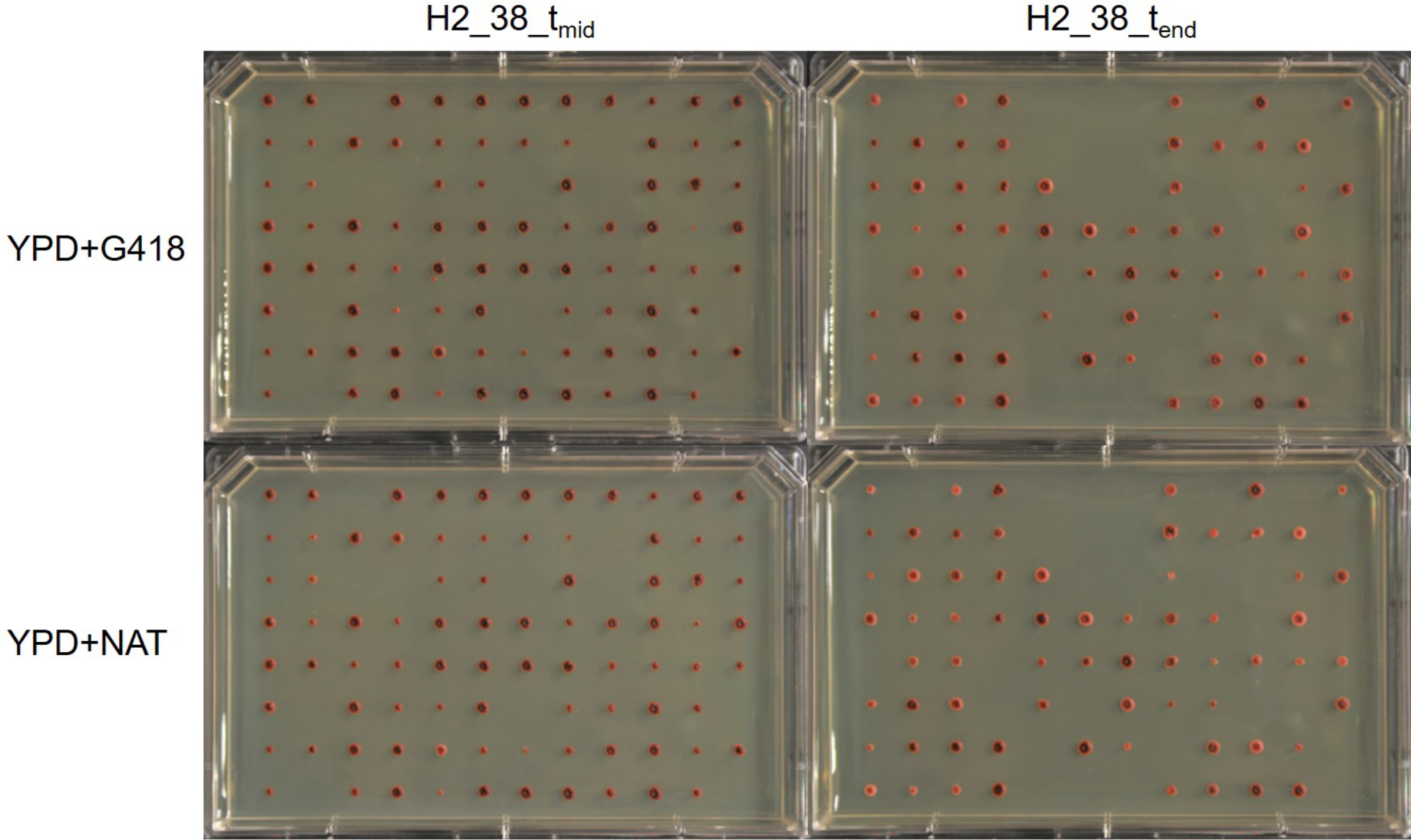
*S. paradoxus* × *S. cerevisiae* tetraploid genome segregates a copy of each parental genome in its diploid spores. The tetraploid line H2_38 spores were plated on selective media to assess the segregation of the selection cassettes during meiosis. S. cerevisiae genome harbor the NAT resistance while S. paradoxus genome harbor the G418 resistance at the HO locus (Chr IV). As all the viable spores inherited both resistances, it is likely that the spores contain full non-recombined *S. cerevisiae* and a *S. paradoxus* haplotypes.

**Table S1.** List of strains used in this study.

| Strain | Sampling site | Lineage | Mating type | Genotype | Reference Wild | Reference hoΔ | Reference ade2Δ |
| --- | --- | --- | --- | --- | --- | --- | --- |
| YPS644 | Buck Hill Falls, PE | *SpA* | $\alpha$ | hoΔ::KAN, ade2Δ::HPH | Kuehne *et al.* 2007 | Charron *et al.* 2014 | This study |
| YPS644 | Buck Hill Falls, PE | *SpA* | a | hoΔ::KAN, ade2Δ::HPH | Kuehne *et al.* 2007 | Charron *et al.* 2014 | This study |
| YPS744 | Tuscarora Forest, PE | *SpA* | $\alpha$ | hoΔ::NAT, ade2Δ::HPH | Kuehne *et al.* 2007 | Charron *et al.* 2014 | This study |
| YPS744 | Tuscarora Forest, PE | *SpA* | a | hoΔ::NAT, ade2Δ::HPH | Kuehne *et al.* 2007 | Charron *et al.* 2014 | This study |
| UWOPS_91_202 | Long Point, ON | *SpB* | a | hoΔ::KAN, ade2Δ::HPH | Kuehne *et al.* 2007 | Charron *et al.* 2014 | This study |
| UWOPS_91_202 | Long Point, ON | *SpB* | a | hoΔ::NAT | Kuehne *et al.* 2007 | Charron *et al.* 2014 | - |
| MSH604 | Mont-St-Hilaire, QC | *SpB* | a | hoΔ::NAT, ade2Δ::HPH | Leducq *et al.* 2014 | Leducq *et al.* 2016 | This study |
| LL12_021 | Pointe Platon, QC | *SpB* | $\alpha$ | hoΔ::NAT, ade2Δ::HPH | Leducq *et al.* 2014 | Leducq *et al.* 2016 | This study |
| LL12_028 | Sherbrooke, QC | *SpB* | $\alpha$ | hoΔ::KAN, ade2Δ::HPH | Leducq *et al.* 2014 | Leducq *et al.* 2016 | This study |
| YPS484 | Grand Bend, ON | *SpB* | a | hoΔ::HPH | Kuehne *et al.* 2007 | Charron *et al.* 2014 | - |
| yHKS226 | Plainfield, NH | *SpB* | $\alpha$ | hoΔ::HPH | Sylvester *et al.* 2015 | Leducq *et al.* 2016 | - |
| LL11_004 | Cap Chat, QC | *SpC* | $\alpha$ | hoΔ::KAN, ade2Δ::HPH | Leducq *et al.* 2014 | Charron *et al.* 2014 | This study |
| LL11_009 | St-Michel-du-Squatec, QC | *SpC* | $\alpha$ | hoΔ::NAT, ade2Δ::HPH | Leducq *et al.* 2014 | Charron *et al.* 2014 | This study |
| LL11_009 | St-Michel-du-Squatec, QC | *SpC* | a | hoΔ::NAT, ade2Δ::HPH | Leducq *et al.* 2014 | Charron *et al.* 2014 | This study |
| MSH_587-1 | Mont-St-Hilaire, QC | *SpC* | $\alpha$ | hoΔ::HPH | Leducq *et al.* 2014 | Charron *et al.* 2014 | - |
| YPS667 | Buck Hill Falls, PE | *SpC* | $\alpha$ | hoΔ::HPH | Kuehne *et al.* 2007 | Charron *et al.* 2014 | - |
| yHKS225 | Plainfield, NH | *SpC* | a | hoΔ::NAT | Sylvester *et al.* 2015 | Leducq *et al.* 2016 | - |
| LL12_019 | Pointe Platon, QC | *SpC* | a | hoΔ::HPH | Leducq *et al.* 2014 | Leducq *et al.* 2016 | - |
| LL13_040 | Rockport, MA | *Scer* | $\alpha$ | hoΔ::KAN, ade2Δ::HPH | This study | This study | This study |
| LL13_040 | Rockport, MA | *Scer* | a | hoΔ::KAN, ade2Δ::HPH | This study | This study | This study |
| LL13_054 | Rockport, MA | *Scer* | $\alpha$ | hoΔ::NAT, ade2Δ::HPH | This study | This study | This study |
| LL13_054 | Rockport, MA | *Scer* | a | hoΔ::NAT, ade2Δ::HPH | This study | This study | This study |

**Table S2.**
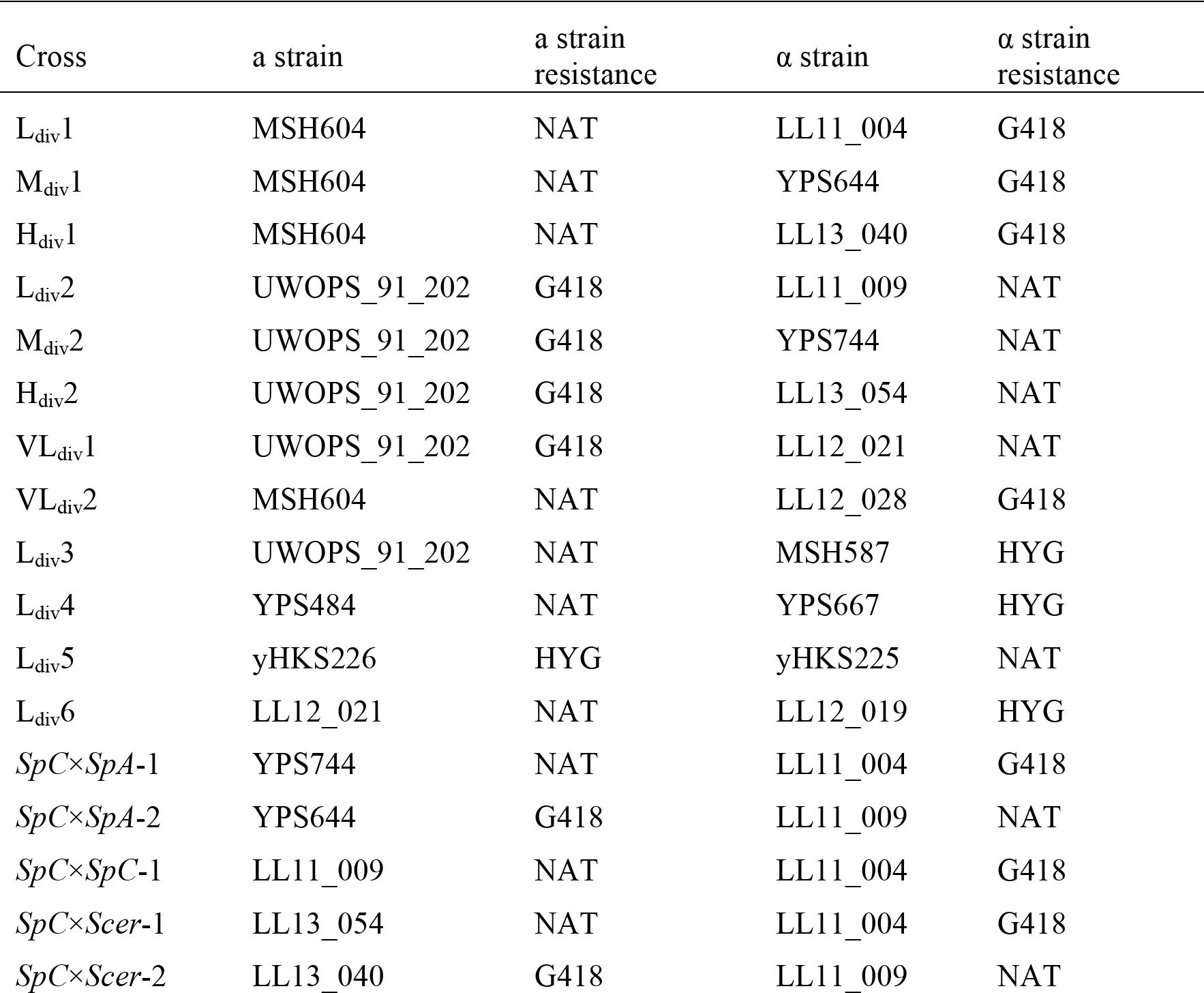
List of crosses used in this study.

| Cross | a strain | a strain<br>resistance | $\alpha$ strain | $\alpha$ strain<br>resistance |
| --- | --- | --- | --- | --- |
| L<sub>div</sub>1 | MSH604 | NAT | LL11_004 | G418 |
| M<sub>div</sub>1 | MSH604 | NAT | YPS644 | G418 |
| H<sub>div</sub>1 | MSH604 | NAT | LL13_040 | G418 |
| L<sub>div</sub>2 | UWOPS_91_202 | G418 | LL11_009 | NAT |
| M<sub>div</sub>2 | UWOPS_91_202 | G418 | YPS744 | NAT |
| H<sub>div</sub>2 | UWOPS_91_202 | G418 | LL13_054 | NAT |
| VL<sub>div</sub>1 | UWOPS_91_202 | G418 | LL12_021 | NAT |
| VL<sub>div</sub>2 | MSH604 | NAT | LL12_028 | G418 |
| L<sub>div</sub>3 | UWOPS_91_202 | NAT | MSH587 | HYG |
| L<sub>div</sub>4 | YPS484 | NAT | YPS667 | HYG |
| L<sub>div</sub>5 | yHKS226 | HYG | yHKS225 | NAT |
| L<sub>div</sub>6 | LL12_021 | NAT | LL12_019 | HYG |
| *SpC*×*SpA*-1 | YPS744 | NAT | LL11_004 | G418 |
| *SpC*×*SpA*-2 | YPS644 | G418 | LL11_009 | NAT |
| *SpC*×*SpC*-1 | LL11_009 | NAT | LL11_004 | G418 |
| *SpC*×*Scer*-1 | LL13_054 | NAT | LL11_004 | G418 |
| *SpC*×*Scer*-2 | LL13_040 | G418 | LL11_009 | NAT |

**Table S3.**
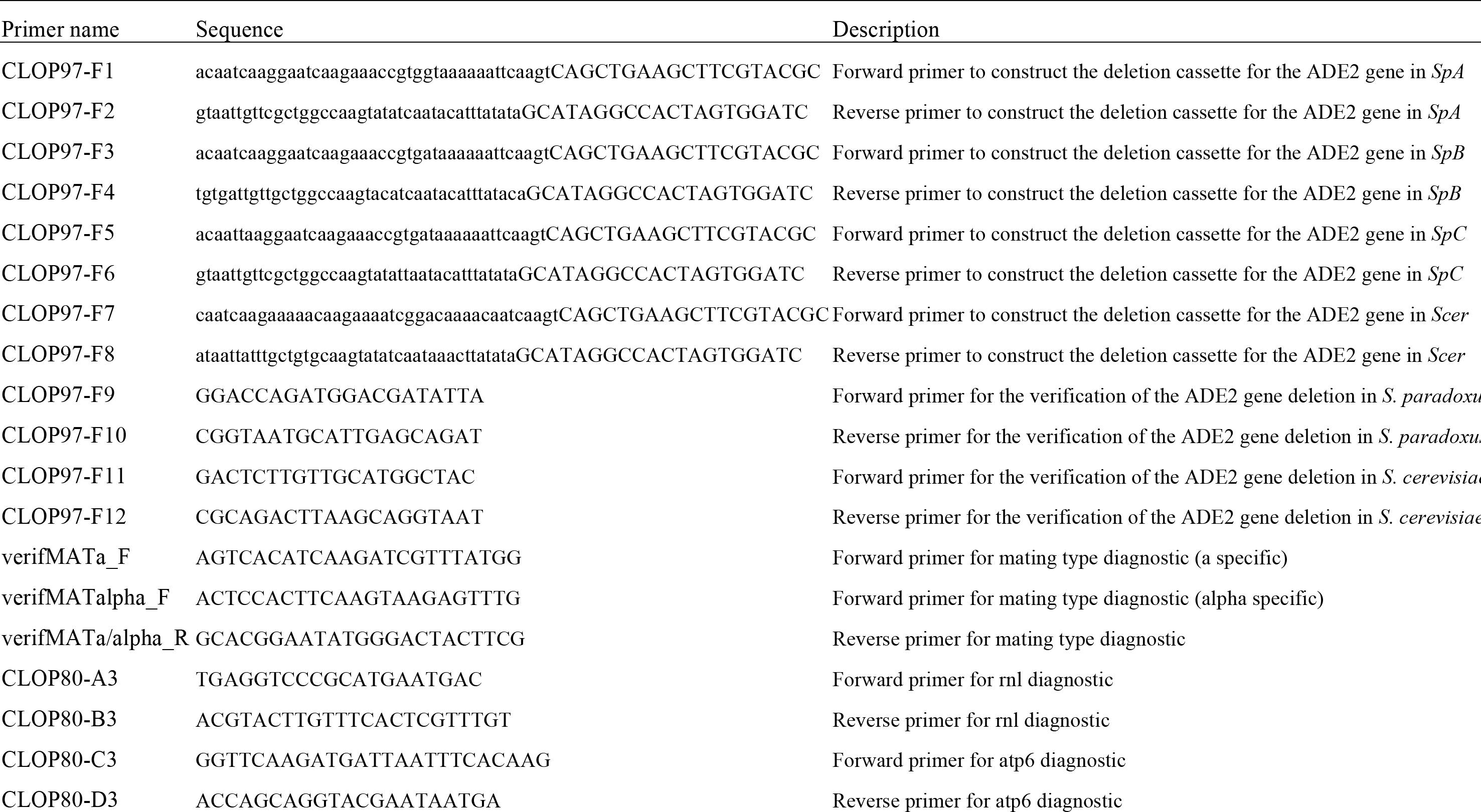
Oligonucleotides used in this study.

**Table S4.** Logrank test P-values for all survival curves.

| Crosses |  |  |  |  |  |  |  |
| --- | --- | --- | --- | --- | --- | --- | --- |
|  | L1 | M1 | H1 | L2 | M2 | H2 | VL1 |
| M1 | 0.147232 | NA | NA | NA | NA | NA | NA |
| H1 | 0.807183 | 0.215188 | NA | NA | NA | NA | NA |
| L2 | 0.630827 | 0.350038 | 0.754435 | NA | NA | NA | NA |
| M2 | 0.00046 | 0.020101 | 0.000528 | 0.000759 | NA | NA | NA |
| H2 | 0.419209 | 0.477771 | 0.599259 | 0.754435 | 0.00108 | NA | NA |
| VL1 | 0.001695 | 0.045283 | 0.002761 | 0.005074 | 0.754435 | 0.007415 | NA |
| VL2 | 0.000528 | 0.006983 | 0.000616 | 0.000895 | 0.235393 | 0.00108 | 0.419209 |

| Cross type |  |  |  |
| --- | --- | --- | --- |
|  | M | VL | L |
| VL | 0.010114 | NA | NA |
| L | 0.000305 | 2.49E-06 | NA |
| H | 0.000786 | 3.45E-06 | 0.667319 |

| Blocks |  |  |  |  |  |
| --- | --- | --- | --- | --- | --- |
|  | set1(1-64) | set1(65-96) | set2(1-64) | set2(65-96) | VL1 |
| set1(65-96) | 0.075385 | NA | NA | NA | NA |
| set2(1-64) | 0.018042 | 0.907645 | NA | NA | NA |
| set2(65-96) | 0.147398 | 0.768516 | 0.689276 | NA | NA |
| VL1 | 0.003756 | 0.042042 | 0.042042 | 0.025266 | NA |
| VL2 | 0.00077 | 0.009335 | 0.009335 | 0.006103 | 0.388572 |

| Groups |  |
| --- | --- |
|  | LH |
| VLM | 5.22E-09 |

**Table S5.** Ploidy distribution of 94 independent hybrids in three replicates of the L_div_1 and L_div_2 crosses.

| Cross | Ploidy |  |  |
| --- | --- | --- | --- |
|  | 2n (%) | 3n (%) | 4n (%) |
| $L_{div1-1}^*$ | 48.0 | 52.0 | 0.0 |
| $L_{div1-2}$ | 98.0 | 2.0 | 0.0 |
| $L_{div1-3}$ | 67.0 | 32.0 | 1.0 |
| $L_{div2-1}^*$ | 42.5 | 57.5 | 0.0 |
| $L_{div2-2}$ | 91.5 | 8.5 | 0.0 |
| $L_{div2-3}$ | 99.0 | 1.0 | 0.0 |

\* crosses used for the evolution experiment

## Data S1. (separate file)

Zip file containing data for fertility (evolution, ITC and autodiploidization), generation number per passage, survival of the strains used in this study and the custom R script to generate Figures.

## Data S2. (separate file)

Raw cytometer data from the ploidy experiments can be downloaded at: https://www.dropbox.com/s/21ak44rqfcut4pk/Data_S2_Cytometry_Data_and_Scripts.zip?dl=0

